# Connexin 46 and connexin 50 gap junction channel open stability and unitary conductance are shaped by structural and dynamic features of their N-terminal domains

**DOI:** 10.1101/2020.07.01.182584

**Authors:** Benny Yue, Bassam G. Haddad, Umair Khan, Honghong Chen, Mena Atalla, Ze Zhang, Daniel M. Zuckerman, Steve L. Reichow, Donglin Bai

## Abstract

The connexins form intercellular communication channels, known as gap junctions (GJs), that facilitate diverse physiological roles in vertebrate species, ranging from electrical coupling and long-range chemical signaling, to coordinating development and nutrient exchange. GJs formed by different connexins are expressed throughout the body and harbor unique channel properties that have not been fully defined mechanistically. Recent structural studies have implicated the amino-terminal (NT) domain as contributing to isoform-specific functional differences that exist between the lens connexins, Cx50 and Cx46. To better understand the structural and functional differences in the two closely related, yet functionally distinct GJs, we constructed models corresponding to CryoEM-based structures of the wildtype Cx50 and Cx46 GJs, NT domain swapped chimeras (Cx46-50NT and Cx50-46NT), and point variants at the 9^th^ residue (Cx46-R9N and Cx50-N9R) for comparative MD simulation and electrophysiology studies. All of these constructs formed functional GJ channels, except Cx46-50NT, which correlated with increased dynamical behavior (instability) of the NT domain observed by MD simulation. Single channel conductance (γ_j_) also correlated well with free-energy landscapes predicted by MD, where γ_j_ of Cx46-R9N was increased from Cx46 and the γ_j_s of Cx50-46NT and Cx50-N9R was decreased from Cx50, but to a surprisingly greater degree. Additionally, we observed significant effects on transjunctional voltage-dependent gating (V_j_-gating) and open-state dwell times induced by the designed NT domain variants. Together, these studies indicate that the NT domains of Cx46 and Cx50 play an important role in defining channel properties related to open-state stability and single channel conductance.

## Introduction

Gap junctions (GJs) are a class of membrane channels that provide a direct passageway between neighboring cells and facilitate the exchange of ions and small molecules (Saez *et al*., 2003; Goodenough & Paul, 2009). This type of cell-to-cell communication is facilitated by a unique channel architecture, whereby a continuous water-filled pore of ~1.5 nm in diameter is formed between two opposing cell membranes, effectively coupling the cytoplasms of adjoined cells (Sosinsky & Nicholson, 2005). These direct passageways are essential for enabling fast transmission of electrical signals in the brain and heart and for facilitating long-range metabolic coupling in most tissues. Because of their important physiological roles, genetic mutations or pathological conditions that lead to aberrant channel function have been linked to a variety of human disease, including deafness, cataracts, peripheral neuropathy, cardiac arrhythmia, stroke, skin disorders, and cancers (Aasen *et al*., 2016; Garcia *et al*., 2016; Delmar *et al*., 2017).

GJ intercellular channels are formed through the assembly of twelve integral membrane proteins called connexins. Six connexins oligomerize to form a hemichannel (also known as connexon) and two hemichannels from neighboring cells can dock together to form a functional gap junction channel if they are docking compatible (White *et al*., 1994b; Bai *et al*., 2018). Connexins are ubiquitously expressed throughout the body, with each connexin displaying tissue specific distribution (Saez *et al*., 2003; Goodenough & Paul, 2009). Humans express 21 connexin isoforms in a cell-type specific fashion, possibly reflecting the need for unique channel functions that match the physiological demands of their environment. Adding to this diversity, different tissues often express more than one type of connexin, which may co-assemble into so-called heteromeric channels (mixed within the same hemichannel) and/or heterotypic channels (mixed between opposing hemichannels) (Koval *et al*., 2014; Bai *et al*., 2018). In this way, cells may fine-tune their GJ channel properties to control the synchronization of physiological activities or to maintain homeostasis.

It has been well-established that GJs formed by different connexins display different channel properties, including distinct permeability to substrates, various rates of ion permeation, and gating control by a variety of factors, including intracellular protons, divalent cations, and transjunctional voltage (V_j_) (Bukauskas & Verselis, 2004; Bargiello & Brink, 2009). Transjunctional voltage dependent gating (also known as V_j_-gating) exists in all characterized GJs, and depending on the component connexins the resultant GJs could show various levels of gating extent, half deactivation voltage, gating charge (which determines gating sensitivity), and gating kinetics (Harris *et al*., 1981; Paul *et al*., 1991; Veenstra *et al*., 1994; Verselis *et al*., 1994; White *et al*., 1994a; Trexler *et al*., 1996; Oh *et al*., 1999; Musa *et al*., 2004). Some of these macroscopic V_j_-gating properties could be visualized at the single GJ channel level, providing mechanistic insights. Different connexin GJs also showed drastically different rates of ion permeation measured by single channel conductance, from a few pico-Siemens (pS) of Cx30.2 and Cx36 to 200 - 300 pS of Cx50 and Cx37 (Veenstra *et al*., 1994; Srinivas *et al*., 1999; Moreno *et al*., 2005; Bukauskas *et al*., 2006). Although such distinct channel functions are well-established, the underlying molecular and structural mechanisms for controlling GJ channel gating and ion permeation have not yet been fully defined.

Structurally, all connexins are predicted to have the same topological structure, which includes a transmembrane domain of four alternating a-helices (TM1-4), two extracellular loops (EC1 and EC2), an amino-terminal (NT) domain, a carboxyl terminus (CT), and a cytoplasmic loop (CL) (Sohl & Willecke, 2004). The first high-resolution structure of a GJ channel was reported for Cx26, using x-ray crystallography (Maeda *et al*., 2009). This structure defined the pore lining domains as TM1, TM2 and EC1, with the NT domain folding into the vestibule at the channel entrance. In addition to other domains, the NT domain and residues within this domain of Cx26, Cx32, Cx40, Cx45, Cx46, and Cx50 have been shown to be important for both V_j_-gating and unitary channel conductance (Verselis *et al*., 1994; Purnick *et al*., 2000; Tong *et al*., 2004; Tong & Ebihara, 2006; Xin *et al*., 2010; Xin *et al*., 2012a). Indeed, the location of the NT domain at the cytoplasmic vestibule is ideally positioned to play a role in multiple functions, including acting as a selectivity filter, sensing transjunctional voltage, and channel gating.

Using single particle electron cryo-microscopy (CryoEM), we recently reported the high-resolution structures of native Cx46 and Cx50 GJ channels, isolated from the mammalian (sheep) lens, at a resolution of 3.4 Å (Myers *et al*., 2018). Cx46 and Cx50 play essential roles in lens development, and in supporting lens transparency by facilitating ionic currents that drive fluid and metabolite circulation in this avascular organ (Mathias *et al*., 2010). In the CryoEM-based structures, the NT domain in Cx46 and Cx50 was shown to adopt a canonical amphipathic helix, with the hydrophobic face buried against TM1/2 and the hydrophilic residues facing the lumen of the channel. This conformation was distinct from that of Cx26, and molecular dynamics (MD) studies revealed that the NT domains of Cx46 and Cx50 adopted a stabilized open-state conformation compared to that of the Cx26 structure and likely establish the selectivity filter and primary energy barriers for ion permeation (Myers *et al*., 2018). Specifically, MD simulations and potential of mean force (PMF) calculations conducted on Cx46 and Cx50 GJs revealed much more favorable free energy pathways to permeation of cation versus anion permeation (Myers *et al*., 2018), a property that has been demonstrated experimentally for rodent Cx50 and Cx46 GJs (Trexler *et al*., 1996; Srinivas *et al*., 1999; Sakai *et al*., 2003; Tong *et al*., 2014). This comparative analysis further revealed a significant free energy barrier for K^+^ permeation at the Cx46 NT domain, which was absent in Cx50, and appeared to be centered around the 9^th^ position, where a positively charged and bulky arginine (R9) residue exists in Cx46.

Based on these observations, we hypothesized that differences in the V_j_-gating and single channel conductance (γ_j_) of Cx46 and Cx50 are controlled in part by their NT domain, especially the 9^th^ residue, where γ_j_ of Cx50 has been shown to be significantly larger than Cx46 in human and rodent GJs (Srinivas *et al*., 1999; Hopperstad *et al*., 2000; Trexler *et al*., 2000; Sakai *et al*., 2003; Xin *et al*., 2010; Rubinos *et al*., 2014; Abrams *et al*., 2018). To better define the key structural and dynamic features that distinguish these channel properties and to more closely align our structure models with functional studies, we employed a combination of MD simulation studies together with dual patch clamp technique to characterize GJ channel properties of sheep Cx46, Cx50, NT domain swapped chimeras (Cx46-50NT and Cx50-46NT), and single point variants at the 9^th^ residue (Cx46-R9N and Cx50-N9R). Our results showed remarkable correlation between our atomic models and observed functional features, which indicate that key differences in the structures of the NT domain and the 9^th^ residue of Cx46 and Cx50 play important roles in defining GJ channel formation, V_j_-gating properties, the stability of the open-state, and in tuning single channel ion conductance properties. Together, these studies provide new insights into the mechanistic principles governing isoform-specific differences in GJ channel properties, and demonstrate that sheep Cx46 and Cx50 provide an ideal system for correlative structure-function studies of GJs.

## Methods

### Plasmid construction

Sheep Cx46 (Cx46, also known as Cx44) and Cx50 (Cx50, also known as Cx49) cDNA were synthesized and each of them was inserted into an expression vector, pIRES2-EGFP, with an untagged GFP reporter between the restriction enzyme sites, XhoI and EcoRI (NorClone Biotech Laboratories, London, Ontario). Cx46-IRES-GFP was used as a template for polymerase chain reaction cloning to generate the chimera, Cx46-50NT, in which the amino terminal (NT) domain of Cx46 was replaced by Cx50 NT domain and a missense variant Cx46-R9N. Similarly, Cx50-IRES-GFP was used as a template to generate Cx50-46NT and Cx50-N9R. The primers used to generate these chimeras and point variants are listed below. Cx46-50NT: forward 5’ ATG GGC GAC TGG AGC TTC CTG GGG AAC ATC TTG GAG GAG GTG AAT GAG CAC TCC ACT GTC ATC 3’ and reverse 5’ GAT GAC AGT GGA GTG CTC ATT CAC CTC CTC CAA GAT GTT CCC CAG GAA GCT CCA GTC GCC CAT 3’ Cx50-46NT: forward 5’ ATG GGA GAC TGG AGT TTC CTG GGG AGA CTC CTA GAG AAC GCC CAG GAG CAC TCC ACG GTC ATC 3’ and reverse 5’ GAT GAC CGT GGA GTG CTC CTG GGC GTT CTC TAG GAG TCT CCC CAG GAA ACT CCA GTC TCC CAT 3’ Cx46-R9N: forward 5’ GAC TGG AGC TTC CTG GGG AAC CTC CTA GAG AAC GCC CAG 3’ and reverse 5’ CTG GGC GTT CTC TAG GAG GTT CCC CAG GAA GCT CCA GTC 3’ Cx50-N9R: forward 5’ GAC TGG AGC TTC CTG GGG AAC CTC CTA GAG AAC GCC CAG 3’ and reverse: 5’ CTG GGC GTT CTC TAG GAG GTT CCC CAG GAA GCT CCA GTC 3’

GFP fusion tagged Cx46 (Cx46-GFP), the chimera (Cx46-50NT-GFP), and point variant (Cx46-R9N-GFP) were generated by subcloning these construct into EGFP expression vector (Kim *et al*., 2013). Mutagenesis was used to remove the stop codon from each of these vectors and ensure that the GFP was linked in frame with a peptide linker (LGILQSTVPRARDPPVAT) between Cx46 (or Cx46-50NT) and GFP.

### Cell culture and transient transfections

Gap junction (GJ) deficient mouse neuroblastoma (N2A) cells (American Type Culture Collection, Manassas, VA, USA) were grown in Dulbecco’s Modified Eagle’s Medium (DMEM) (Life Technologies Corporation, Grand Island, NY, USA) containing 4.5 g/L D-(+)-glucose, 584 mg/L L-glutamine, 110 mg/L sodium pyruvate, 10% fetal bovine serum (FBS), 1% penicillin, and 1% streptomycin, in an incubator with 5% CO2 at 37° C (Sun *et al*., 2013). N2A cells were transfected with 1.0 μg of a cDNA construct and 2 μL of X-tremeGENE HP DNA transfection reagent (Roche Diagnostics GmbH, Indianapolis, IN, USA) in Opti-MEM + GlutaMAX-I medium for 4 hours. After transfection, the medium was changed back to FBS-containing DMEM and incubated overnight. The next day, N2A cells transfected with Cx46 and its variants were replated onto glass coverslips for 2 hours before transferring to the recording chamber. N2A cells transfected with Cx50 and its variants were replated onto glass coverslips for 10 hours before transferring to the recording chamber. Isolated green fluorescent cell pairs were selected for patch clamp study of homotypic GJs.

### Electrophysiological recording

Glass coverslips with transfected cells were placed into a recording chamber on an upright microscope (BX51WI, Olympus). The chamber was filled with extracellular solution (ECS) at room temperature (22 – 24° C). The ECS contained (in mM): 135 NaCl, 2 CsCl, 2 CaCl_2_, 1 MgCl_2_, 1 BaCl_2_, 10 HEPES, 5 KCl, 5 D-(+)-glucose, 2 Sodium pyruvate, pH adjusted to 7.4 with 1M NaOH, and osmolarity of 310-320 mOsm. Dual whole cell patch clamp was performed on green fluorescent cell pairs with MultiClamp 700A amplifier (Molecular Devices, Sunnyvale, CA, USA). Patch pipettes were pulled with a micropipette puller (PC-10, Narishige International USA Inc., Amityville, NY, USA) and filled with intracellular solution (ICS) containing (in mM): 130 CsCl, 10 EGTA, 0.5 CaCl_2_, 5 Na2ATP, 10 HEPES, adjusted to pH 7.2 with 1 M CsOH, and osmolarity of 290-300 mOsm. After establishing whole cell recording on the cell pair, both cells were voltage clamped at 0 mV. In one cell of the pair, a series of voltage pulses (±20 to ±100 mV) were applied to establish transjunctional voltage (V_j_). The other cell of the pair was constantly held at 0 mV to record gap junctional current (I_j_). The current was low-pass filtered (Bessel filter at 1 kHz) and recorded using pClamp9.2 software at a sampling rate of 10 kHz via an AD/DA converter (Digidata 1322A, Molecular Devices, Sunnyvale, CA, USA).

### Transjunctional voltage dependent gating

Transjunctional voltage (V_j_) dependent gating (V_j_-gating) was studied in cell pairs expressing one of the constructs. Voltage pulses (±20 to ±100 mV with 20 mV increment) were delivered in one cell of the pair to establish V_j_s and transjunctional currents (I_j_s) were recorded in the other cell. In most cases, I_j_s peaked at the beginning and then deactivated (especially with high V_j_s, ±40 to ±100 mV) to a steady-state near the end of a 7 second V_j_ pulse. Gap junctional conductance (G_j_) was calculated (G_j_ = I_j_/V_j_). The steady state G_j_ was normalized to the peak G_j_ to obtain a normalized steady-state junctional conductance (G_j,ss_) for each tested V_j_s. The G_j,ss_ were then plotted with V_j_s to obtain a G_j,ss_ – V_j_ plot, which could sometimes fit well with a two-state Boltzmann equation for each V_j_ polarity to obtain gating parameters, V_0_, G_min_, G_max_, and *A* (Jassim *et al*., 2016). V_0_ is the voltage when the G_j,ss_ is reduced by half [(G_max_ - G_min_)/2], G_min_ is the normalized minimum residual conductance, while G_max_ represents the maximum normalized conductance, and *A* is the slope of the curve which reflects V_j_-gating sensitivity (Spray *et al*., 1981).

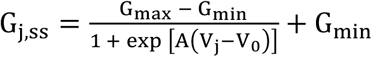

### Single Channel Analysis

Single channel current (i_j_) could be observed in cell pairs with few active GJ channels. The recorded currents were further filtered using a low-pass Gaussian filter at 200 Hz in Clampfit9.2 for measuring current amplitude and display in figures. The amplitude i_j_s for fully open state at different V_j_s were measured with fitting Gaussian functions on all point current amplitude histograms. The i_j_s of different cell pairs were averaged under the same V_j_, regardless of V_j_ polarity, to generate i_j_ – V_j_ plot. Linear regression of i_j_ – V_j_ plot with at least 4 different V_j_s was used to estimate slope unitary GJ channel conductance (also known as single channel conductance, γ_j_).

Single GJ channel open dwell times were analyzed on cell pairs with only one active GJ channel. In this case, single channel currents were filtered with a low-pass Gaussian filter at 500 Hz (Clampfit9.2). The open dwell time was measured at the V_j_s from ±40 to ±100 mV using Clampfit9.2. Any single channel open events with less than 2 ms were excluded from the analysis as this is beyond the resolution of our single channel recordings. The average open dwell time at each V_j_ was calculated and plotted for comparison. The total number of open events for Cx50 and Cx50-N9R GJs was a lot higher than other GJs and suitable for more in depth analysis on the open dwell times. Histograms of Cx50 and Cx50-N9R were plotted in a logarithmic scale with 5 bins/decade and fitted with a two-exponential probability fitting. The two time constant (τ) values represent the corresponding open dwell times for each of the peaks in the fitting. The τ_mean_ value was calculated by taking the weighted average of the two τ values using the area under each peak.

### Molecular dynamic simulations

Visual Molecular Dynamics (VMD) v.1.9.3 was used to build the dodecameric channels for Cx46 (PDB: 6MHQ) and Cx50 (PDB: 6MHY) wild-type systems for molecular dynamics (MD) simulations (Humphrey *et al*., 1996; Myers *et al*., 2018). Sidechains were protonated according to neutral conditions, and the protonated HSD model was used for all histidine residues. Disulfide bonds identified in the experimental structures were enforced for both models. Amino acids corresponding to the cytoplasmic loop (CL) connecting TM2–TM3, and the C-terminal (CT) domain of Cx46 and Cx50 were not included for molecular dynamics simulation, as experimental data describing the structure of these large domains (~50 residue CL and ~200 residue CT domain in Cx46 and Cx50) are missing. The introduced N- and C-terminal residues resulting from the missing CL segment (L97 and L142 in Cx46; V97 and L154 in Cx50) were neutralized. NT acetylation sites were introduced in VMD through an all-atom acetylation patch using the AutoPSF plugin, to mimic the *in vivo* co-translational modification identified in the native proteins (Myers *et al*., 2018). The prepared protein structures were submerged in a hydration shell using Solvate v.1.0.177. Water was removed from sections of the channels corresponding to transmembrane domains, based on hydrophobic character and localization of amphipol observed in experimental CryoEM data (~20–50 Å from the center of the channel). To mimic a cell-to-cell junction, the VMD Membrane Builder plugin was used to add two lipid bilayers to each system, containing 1-palmitoyl-2-oleoyl-sn-glycero-3-phosphocholine (POPC), with dimensions of 152 × 152 Å.

The four structural models of designed variants, Cx46-R9N, Cx46-50NT, Cx50-N9R and Cx50-46NT, were also built using VMD, as follows. First, protein-only structures of the respective wildtype models were mutated, as per the sequence differences of the NT domain (defined as residue 2 – 19) for the NT domain chimeras or just the 9^th^ residue for the point variants, using VMD’s *mutator* plugin (Humphrey *et al*., 1996). The variant models were then merged with the lipid bilayers created for their respective wildtype models, described above, using VMD’s *mergestructs* plugin. Lipids which overlapped with the protein models were then removed. Each model was then placed in a water box with dimensions 150 × 150 × 180 Å using VMD’s *solvate* plugin. The models were neutralized using the Autoionize plugin, then 150 mM KCl and 150 mM NaCl were added to the solvent areas corresponding to intracellular and extracellular regions of the simulation box, respectively.

GPU-accelerated Nanoscale Molecular Dynamics v.2.13 (Phillips *et al*., 2005) was used for all classical molecular dynamics (MD) simulations, using the CHARMM36 force field (Huang & MacKerell, 2013) for all atoms and TIP3P explicit model for water. Each model was prepared following the same minimization and equilibration protocol, as follows. First, the lipid tails were allowed to minimize with all other atoms fixed for 1 ns using a 1 fs timestep, allowing the acyl chains to “melt” with constant volume at a temperature of 300 K (NVT). All subsequent simulations were performed using the Langevin piston Nose-Hoover method for pressure control (NPT). Next, the entire system, including lipids, solvent and ions, was allowed to minimize with the protein harmonically constrained (1 kcal mol^−1^), for 1 ns using a 1 fs timestep. A third 1 ns minimization step was then applied, in which the system was free to minimize with a harmonic constraint applied only to the protein backbone (1 kcal mol^−1^), to ensure stable quaternary structure while sidechains relax in their local environment. The entire system was then released from all constraints and subject to all-atom equilibration using a Langevin thermostat (damping coefficient of 1 ps^−1^), with a constant temperature of 310 K and constant pressure of 1 atm, using a 2 fs timestep for 30 ns. Periodic boundary conditions were used to allow for the particle mesh Ewald calculation of electrostatics. Finally, all models were simulated for 100 ns of production, split into four 25 ns replicas to decorrelate simulations and enhance sampling in the local conformational space, where each replica started from the end of 30 ns equilibration with velocities reinitialized, using a 2 fs timestep. A table summarizing the simulation conditions are provided in Supplemental Table 1.

Root mean squared deviations (r.m.s.d.), comparing the backbone conformations of MD simulation to the original starting structures, and root mean square fluctuations (r.m.s.f.) comparing the amplitudes of backbone fluctuations during MD simulation, were calculated using VMD (Supplemental Fig. 1). All six GJ models approached a steady r.m.s.d. during the first 20 ns of the equilibration phase and maintained stability during all production runs.

Population distribution functions were constructed by monitoring the distance between these residues at the 9^th^ and 12^th^ position in adjacent (i → i+1) subunits (in the clockwise direction when viewed from the cytoplasmic vestibule). Distances between functional groups were recorded at every tenth step of the trajectory across all four production runs, for Cx50, Cx46 and each of the variants. The points of reference used to measure the interatomic distance were chosen to capture equivalent rotameric states, as follows: C_ζ_ for R9, C_γ_ for N9, and C_δ_ for E12. Histograms (bin size = 0.1 Å) were normalized and plotted as probability density functions.

### Energetics and analysis of ion permeation pathway

Potentials of mean force (PMF), or the energy landscape, describing the permeation of K^+^ and Cl^-^ were calculated for all systems (Supplemental Fig. 3). To calculate the PMF, a Markov State Model (MSM) was constructed by defining state-space as the position of an ion along the pore-axis (z-axis), which was subdivided into 3 Å bins. A transition matrix (T), which describes the time evolution of the system, was constructed from the conditional probabilities T_ij_ of an ion ending up in state j after a given lag-time, having begun in state i. As in previous work, we employed a short lag-time (2 ps), ensuring the vast majority of transition probabilities occurred between nearest neighbors (*i.e*., i-1 ↔ i ↔i+1). The principle of detailed balance guarantees that any connected paring of states (e.g., neighbors only) is sufficient to determine the unique equilibrium distribution independent of lag time. Transition probabilities can be approximated by counting the instances of transitions at every lag time and storing the values in a transition count matrix. The count matrix was then row normalized to achieve an approximate transition probability matrix (T)

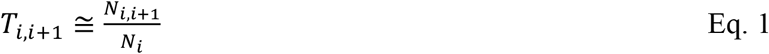

Where *N*_*i,i*+1_ is the number of transitions from state (*i*) to state (*i+1*) in a lag time *τ*, and *N_i_* are the number of times an ion was found in state *i*. The thermodynamics underlying ionic permeation may then be extracted using the principle of detailed balance (*i.e*. statistical equilibrium) (Eq. 2) and Boltzmann statistics (Eq. 3).

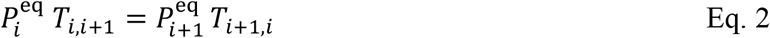

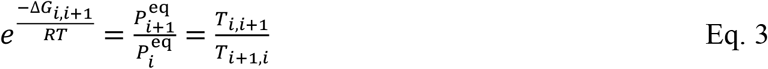

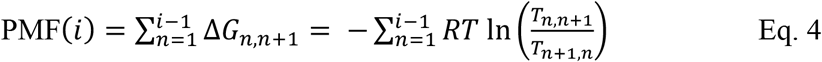

Here, 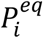 is the probability that an ion will be in each respective bin once equilibrium is achieved, Δ*G*_*i,i*+1_ is the free energy difference between the two states, R is the ideal gas constant (1.986 cal mol^−1^ K^−1^), and T is temperature (310 K) (Eq. 4). Values from the PMF were mapped to the z-axis, interpolated and smoothed using a b-spline. Final PMFs were symmetrized around the center of the channel and adjusted such that the bulk regions were at zero. To enable sufficient sampling of Cl^-^ ions in the channel, a distributed seeding approach was implemented where an individual Cl^-^ is randomly placed within the channel, followed by short 10 ns simulations, and repeated until sufficient sampling for the MSM is achieved. Further explanation and a detailed justification of the distributed seeding approach, and PMF calculation can be found in (Myers *et al*., 2018).

Coulombic surface potentials were calculated using the Adaptive Poisson Boltzmann Solver (APBS) (Jurrus *et al*., 2018) within Chimera(Pettersen *et al*., 2004), using standard settings. Pore profile analysis or the radius at each point along the pore-axis (Z-axis) was calculated using the program HOLE (Smart *et al*., 1996). The calculation is done by rolling a sphere with the radius of a water molecule over the Van der Waals surface of the pore, with the beginning/end of the pore being defined as having a maximum radius of 12 Å. To assess the average pore profile of each model obtained by MD simulation, a snapshot of the protein was saved every 1000th frame (i.e. 2 ns), symmetrized, and then averaged together to provide the average pore profile with the standard error of the mean (SEM) (Supplemental Fig. 2).

### Statistical Analysis

Data are expressed as mean ± SEM. Kruskal Wallis followed by a Dunn’s post-hoc test was used to compare multiple groups of non-parametric data, as specified. One-way ANOVA was used to compare multiple groups of data with Gaussian distribution. Statistical significance is indicated with the asterisks on the graphs only for biologically meaningful group comparisons (\**P* < 0.05; \*\**P* < 0.01; or \*\*\**P* < 0.001). MD data are expressed as mean ± SEM, with the exception of the r.m.s.f. data, which are expressed as the mean ± the 95% confidence interval. Statistical significance is indicated as above, calculated via a standard two-tailed p-value test.

## Results

### Design of variants to probe channel properties of Cx46 and Cx50

The NT domains of Cx46 and Cx50 adopt an amphipathic helix that folds into the GJ pore, contributing to the cytoplasmic vestibules of the channel and differ at only five sites (Fig. 1a,b). Proteomics analysis on native proteins isolated from the lens indicated that Met1 is removed and the resulting NT Gly2 position is acetylated co-translationally in both isoforms (Shearer *et al*., 2008; Wang & Schey, 2009; Myers *et al*., 2018). The first difference in structure occurs at position 9, which is a positively charged arginine in Cx46 and a neutral asparagine in Cx50 (indicated by asterisk in Fig. 1a,b). In the CryoEM-based models, R9 of Cx46 restricts the pore diameter to ~8.8 Å and was identified as forming the main constriction site within the NT domain (Fig. 1c, and (Myers *et al*., 2018)). In comparison, Cx50’s diameter at this site is significantly wider in the CryoEM-based model, at 12.4 Å, (with a minimum constriction of ~11.4 Å located lower on NT domain near S5/D3) (Fig. 1c). Further comparisons show the NT of Cx50 lacks any positively charged amino acids, with an overall net charge of −4, whereas the net charge on the Cx46 NT domain is −2 (Fig. 1b). Other differences localize to the identity of the hydrophobic positions 10 and 14, which form packing interactions with TM1/2 (Fig. 1a,b).

**Fig. 1.**
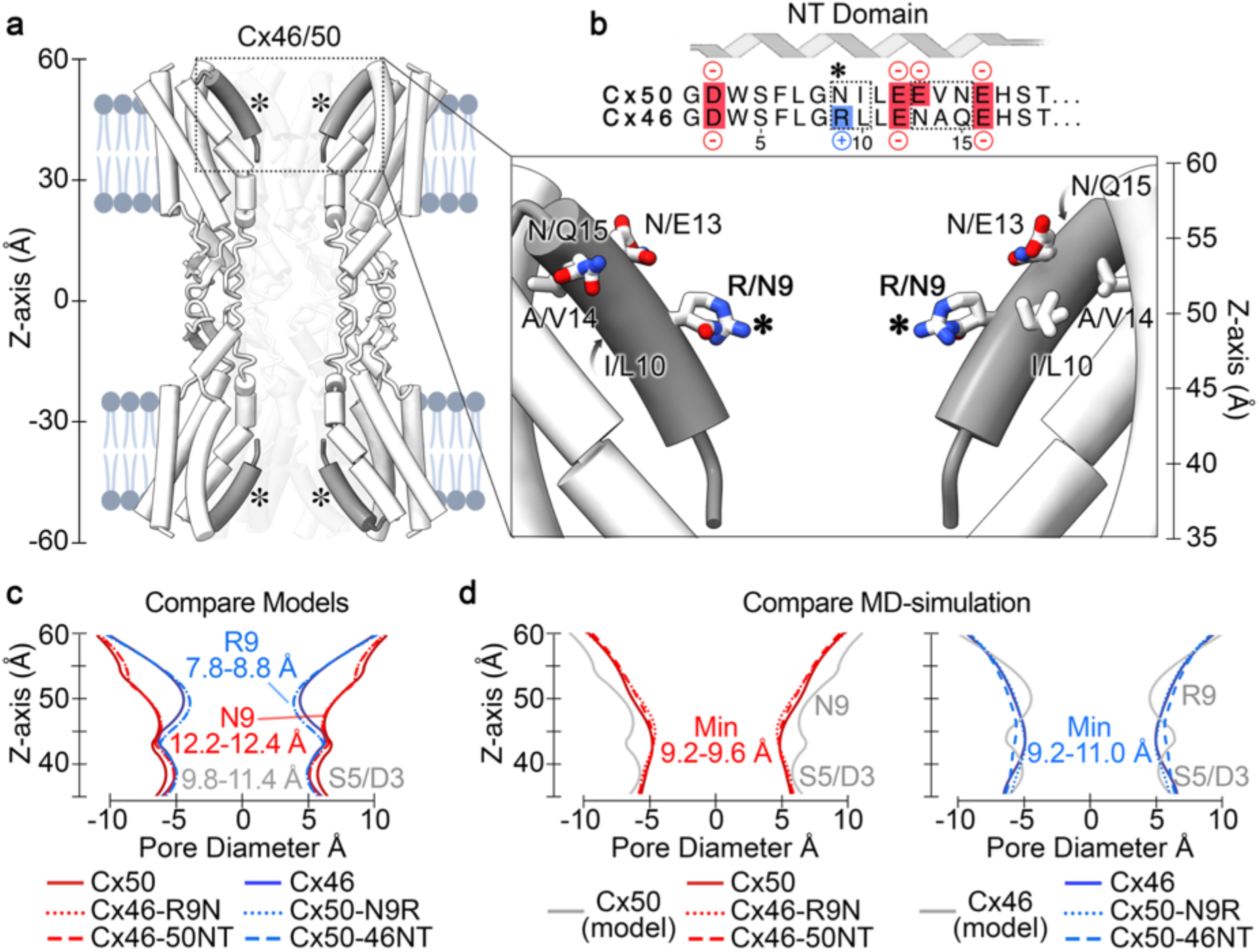
The NT-domain constriction sites of Cx50, Cx46 and designed variants are augmented following MD equilibration. **a.** Cut-away view of Cx46/50 GJ channel shown as tube representation with the NT domain in grey. (*inset*) Shows a zoom view of the NT domain with residue sidechains displayed at sites of genetic variability between Cx50 and Cx46, respectively. (Asterisk, *) indicates location of residue 9 (Arginine in Cx46 and Asparagine in Cx50). **b.** Primary sequence alignment of the Cx50 and Cx46 NT domain (residues 2 – 19). Negatively charged residues are colored in red, positively charged residues are indicated in blue. Sites of variation between the two isoforms are indicated (dotted box). Met1 is not included, as this site is co-translationally removed and Gly2 is acetylated in both isoforms (Myers *et al*., 2018). **c.** Pore diameter analysis of CryoEM based structures of Cx50, Cx46 and the designed NT domain variants. **d.** Averaged pore diameter of Cx50, Cx46 and designed NT domain variants following MD equilibration (pore profile of the Cx46 and Cx50 starting structures are shown for comparison in grey).

Based on these structural comparisons, and previous MD-studies which indicated that Cx46 introduces a significant energetic barrier to K^+^ permeation as compared to Cx50 that is localized to the NT domain (Myers et al 2018), we constructed four models designed to probe the mechanistic and functional role of the NT-domain and the residue identity at the 9^th^ position for comparative all-atom equilibrium MD-simulation and electrophysiology studies in GJ deficient N2A cells. To facilitate the most direct comparisons, all models and constructs used for experimental characterization were based on the sheep Cx46 and Cx50 structures, and correspond to NT domain swapped chimeras (Cx46-50NT and Cx50-46NT), and single point variants at the 9^th^ residue (Cx46-R9N and Cx50-N9R). These models and/or constructs were then used to test the hypothesis that the NT domain and/or position-9 are important in defining key differences in channel properties that exist between the closely related Cx46 and Cx50 GJs.

### Position 9 does not form the primary NT domain constriction site in Cx46, Cx50 or their variants during MD simulation

Constructed models of NT domain swapped chimeras and single point variants at position-9 of Cx46 and Cx50 were built by *in silico* mutation, using VMD (Humphrey *et al*., 1996), and the NT position (G2 in all models) was acetylated (Myers *et al*., 2018). Pore profile analysis, using the program HOLE (Smart *et al*., 1996), show the resulting models produced pore constriction sites as expected based on the identity of position-9 (Fig. 1c), with some differences resulting from the precise conformation selected by the modeling program for the position-9 variants.

Following all-atom MD equilibration in explicit water, the NT domains of all models appeared well behaved and maintained alpha-helical secondary structure (Supplemental Fig. 1). However, the pore profiles of all constructs were modified within the NT domains, as compared to their starting structures, primarily through reorientation of pore-lining sidechain residues. In contrast to the experimental starting models, the ensemble of structures obtained by MD displayed averaged steric landscapes that were all very similar to each other (Fig. 1d and Supplemental Fig. 2). Cx50, Cx46-50NT and Cx46-R9N all converged to a similar profile, with a primary constriction site of ~9.2 – 9.6 Å that is slightly smaller than the experimental model of Cx50, defined primarily by S5/D3 (Z-axis approximately ± 40 Å) (Fig. 1d, red traces). In comparison, Cx46, Cx50-46NT and Cx50-N9R all converged to a similar profile that is somewhat larger than the experimental model of Cx46, ranging between ~9.6 for Cx46 and Cx50-N9R and ~11.0 Å for Cx50-46NT (Fig. 1d, blue traces). The wider diameter of Cx50-46NT is of potential interest to channel function, and discussed in further detail in the following sections.

Notably, in all related models, the R9 position is no longer contributing as the primary constriction site. Upon closer inspection, it is observed that R9 adopts a dynamic exchange of conformational states during MD simulation of Cx46, Cx50-46NT and Cx50-N9R, that reorients the sidechain away from the pore permeation pathway (Fig.2). These conformations include the formation of a salt bridge between the positively charged R9 sidechain and the negatively charged E12 position (heavy atom distance = 2.6 Å) of a neighboring subunit at the (i+1) position (Fig. 2b,c), which is a conserved site in all models (see Fig.1b). Amongst the three variants containing an arginine at position-9, this interaction accounts for ~30-50% of the conformational states observed by MD (Fig. 2e). An alternative, and less populated, conformational state is also observed, where R9 forms an optimal π-π stacking interaction with guanidinium group of a neighboring R9 (planar distance = 3.5 Å) from an adjacent subunit at the i+1 position (Fig. 2b,d) (Vernon *et al*., 2018). This state appears to be further stabilized by salt bridge formation between R9 and E12 of the same subunit (Fig. 2b,d), or between neighboring subunits (similar to the state shown in Fig. 2c). This conformation was observed in ~5-10% of the total conformational states observed by MD (Fig. 2f). In all cases, these stabilizing interactions effectively withdraw the large R9 sidechain away from the permeation pathway and effectively stabilize R9 against the lumen of the channel. Equivalent interactions were not observed in variants where position-9 is occupied by asparagine, with N9 appearing to adopt more random (fluctuating) conformational states (Fig. 2e,f).

**Fig. 2.**
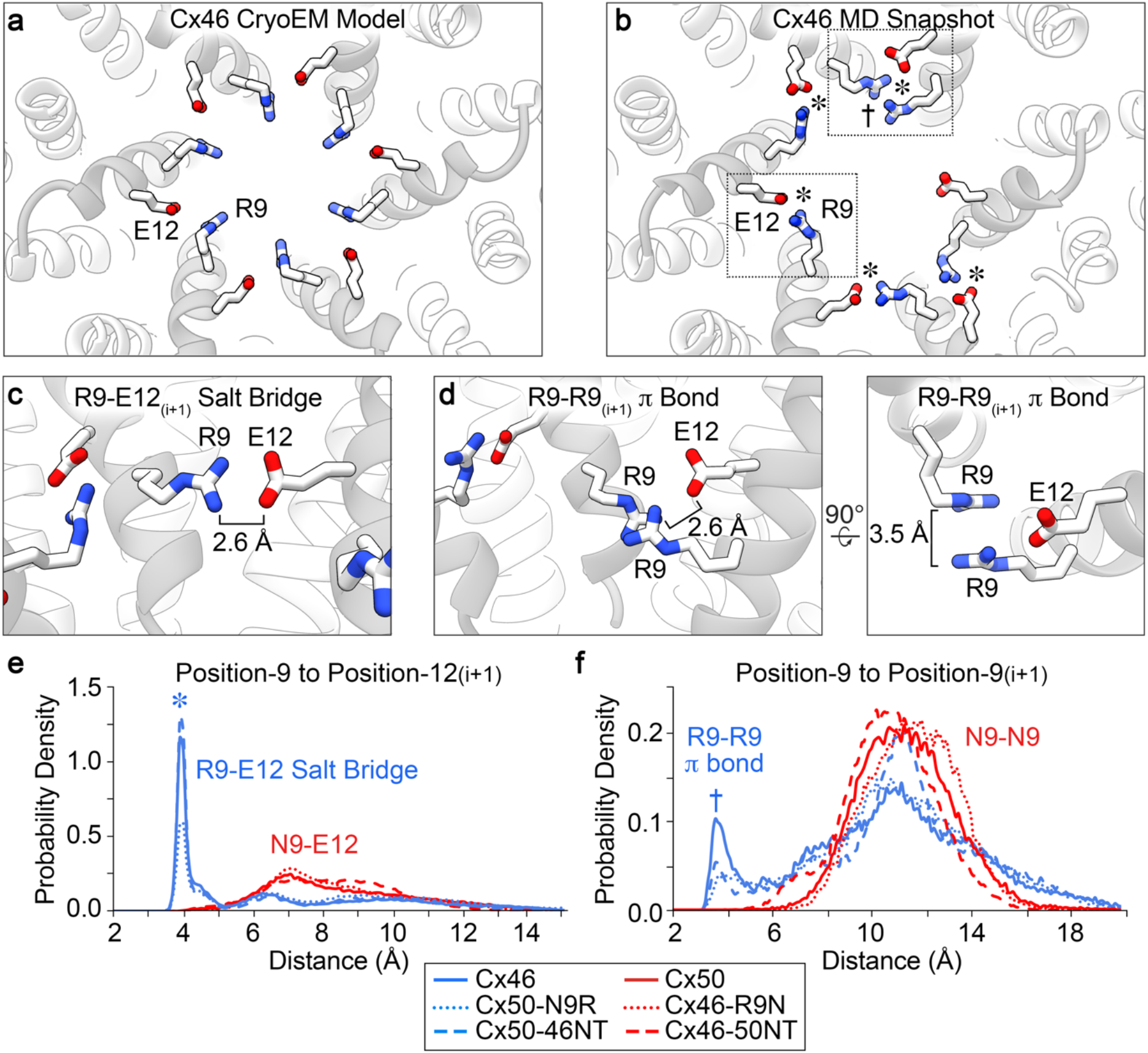
Arginine-9 of Cx46 and designed variants adopts a dynamic ensemble of stable conformational states that prevent pore constriction. **a.** CryoEM-based structure of Cx46 (PDB 6MHQ), viewed along the pore axis with residues R9 and E12 displayed in stick representation. R9 is oriented toward the center of the pore permeation pathway, and forming a primary constriction site. **b.** Representative snap-shot of Cx46 following MD equilibration, viewed as in panel a. R9 adopts alternative conformational states that reorient this sidechain away from the pore permeation pathway, that appear to be stabilized by salt bridge interaction with a neighboring E12 (asterisk, *) and/or π-π interactions with R9 from a neighboring subunit (dagger, †). **c.** Zoom view (boxed in panel b) of a representative R9 salt bridge interaction with a neighboring E12 residue. **d.** Zoom view (boxed in panel b) of a representative π-π interaction between two R9 residues of neighboring subunits. Distances displayed in panels c,d are between nearest heavy atoms. **e.** Probability distribution of the distance between position-9 to position-12 in the neighboring subunit obtained from MD simulation for Cx46, Cx50-N9R and Cx50-46NT (blue traces, represent R9-E12_(i+1)_ distances) and Cx50, Cx46-R9N and Cx46-50NT (red traces, represent N9-E12_(i+1)_ distances). f. Probability distribution of the distance of position-9 to position-9 in the neighboring subunit obtained from MD simulation for Cx46, Cx50-N9R and Cx50-46NT (blue traces, represent R9-R9_(i+1)_ distances) and Cx50, Cx46-R9N and Cx46-50NT (red traces, represent N9-N9_(i+1)_ distances). Distances displayed in panels e,f were measured between R9 C_ζ_ or N9 C_δ_ and E12 C_γ_ to capture equivalent rotameric states, and are therefore greater than those displayed in panels c,d.

The reorganization of the NT domain sidechain conformations observed during MD simulation is consistent with the experimental CryoEM data that was used to build the atomic starting models. This is because, in contrast to the hydrophobic NT anchoring sites, the conformational states of the pore-lining NT domain residues, especially position-9, were not well defined by experimental CryoEM density (presumably reflecting the specimen heterogeneity and/or intrinsic dynamic behavior of these residues) (Myers *et al*., 2018).

### Position 9 defines a key energetic difference in K^+^ ion permeation in Cx50 and Cx46, and their mutational variants

Given the similarities between Cx50 and Cx46 pore constriction sites observed by MD, we next sought to understand how postion-9 and the NT domain may contribute to differences in their respective energetic barriers to ion permeation by evaluating the potential of mean force (PMF) for Cl^-^ and K^+^, which can be extracted from the MD data to describe the free-energy landscapes of these ions along the permeation pathway (*i.e*., the Z-axis) (Fig. 3 and Supplemental Fig. 3). It is noted that inspection of the Coulombic surface of the Cx50 starting model shows that this isoform lacks any positive charge within the pore, when the NT is acetylated (Fig. 3a). In contrast, Cx46 possesses a prominent ring of positive Coulomb potential, belonging to R9, that line the cytoplasmic vestibules at both ends of the channel (Z-axis approximately ± 50 Å) (Fig. 3a). The Coulombic surface potentials of the resulting mutation models are similar to their wild-type counterparts based on the presence or absence of an arginine at the 9^th^ position (not shown).

**Fig. 3.**
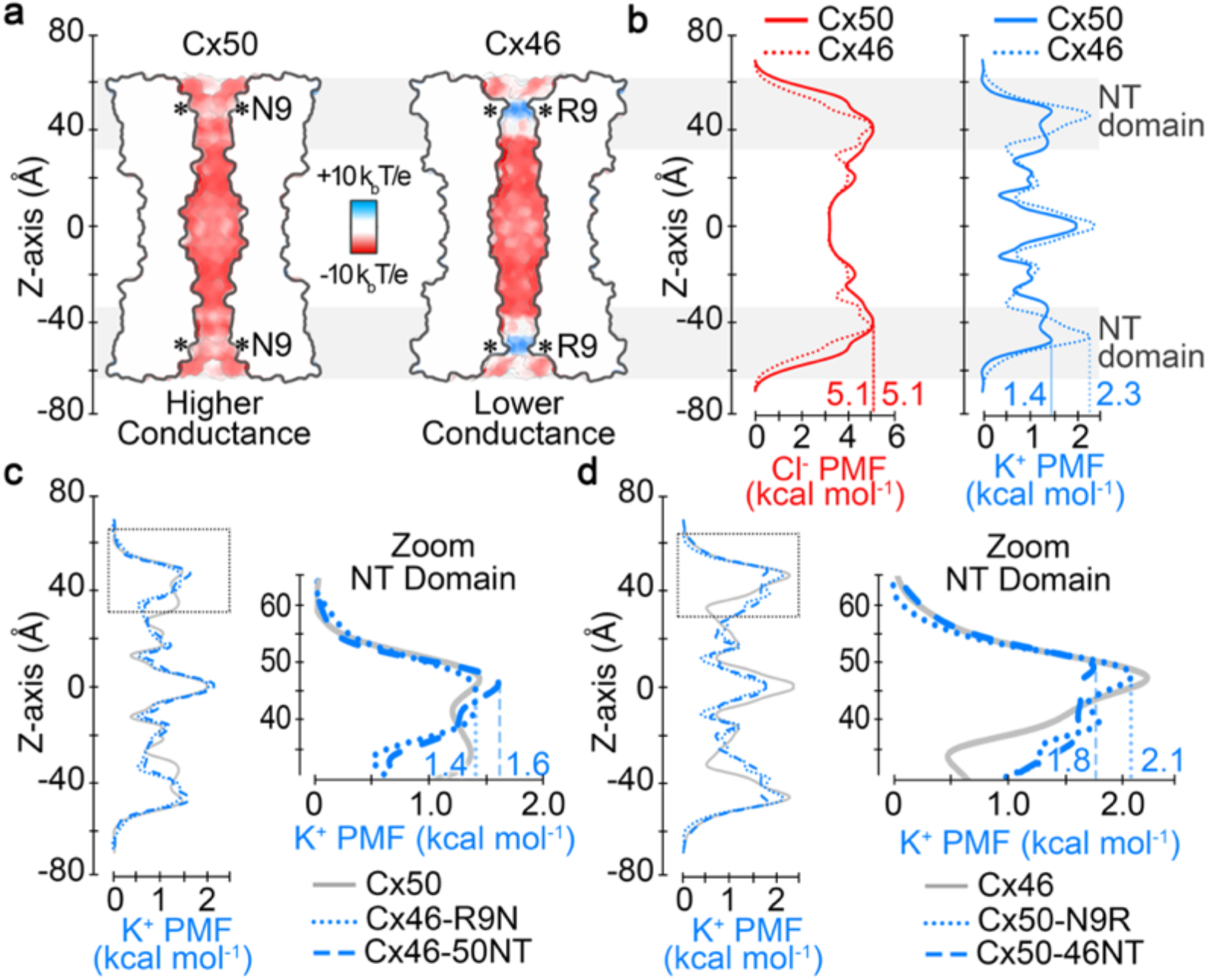
Electrostatics and energetics of the ion permeation pathways of Cx50, Cx46 and designed NT domain variants. **a.** Cut-away surface representation of Cx50 and Cx46, colored by Coulombic potential (red, negative; white, neutral; blue, positive). *T*, temperature; *k*, Boltzmann constant; *e*, charge of an electron. **b.** Potential of mean force (PMF) describing the free-energy landscape (*ΔG*) experienced by Cl^-^ ions (red traces) and K^+^ ions (blue traces) permeating the channel pore of Cx50 and Cx46, as compared to the bulk solvent. **c.** PMFs of K^+^ ions for designed variants Cx46-R9N and Cx46-50NT, with PMF for Cx50 displayed for comparison (gray trace). (*inset*) Shows a zoom view of K^+^ PMFs corresponding to the NT domain in panels c (dotted box region). **d.** PMFs of K^+^ ions for designed variants Cx50-N9R and Cx50-46NT, with corresponding PMF for Cx46 displayed for comparison (gray trace). (*inset*) Shows zoom view of K^+^ PMFs corresponding to the NT domain in panels f (dotted box region). PMFs were symmetrized along Z-axis, and peak energetic barriers identified within the NT domains are indicated. Asterisk (*) in panel a indicates location of position 9. Grey box in panels a,b indicate the region of the NT domain.

Consistent with their Coulombic surface properties, the MD derived PMFs for Cl^-^ showed a high peak energetic barrier, ΔG_Cl-_ ~5.1 kcal mol^−1^ for both Cx50 and Cx46 (defined as the difference between bulk solvent and peak PMF barrier to Cl^-^), which is at least twice as high as the peak energetic barriers to K^+^ (Fig. 3b). The primary energetic barrier to both Cl^-^ and K^+^ are established within the NT-domain region of the channels, and are consistent with this domain acting as the selectivity filter and with the displayed preference for cations that has been previously demonstrated experimentally for both Cx50 and Cx46 (Trexler *et al*., 1996; Srinivas *et al*., 1999; Sakai *et al*., 2003; Tong *et al*., 2014). For these reasons, we have focused the following comparative analysis to the energetic differences between Cx50, Cx46 and the designed variants to their energetics of K^+^ (the major permeant ion). Cl^-^ PMFs for all models were similar and can be found in Supplemental Fig. 3.

The most distinct energetic differences between Cx50 and Cx46 corresponds to the region of the NT domain that aligns with position-9, where Cx46 has a peak energy barrier to K^+^ of ~2.3 kcal mol^−1^ and Cx50 has a peak barrier of ~1.4 kcal mol^−1^, values that agree with our previous study using slightly modified simulation conditions (see Methods) (Myers *et al*., 2018). The difference in peak K^+^ PMF barriers in these two models (ΔΔG_K+_ = 0.9 kcal mol^−1^) is expected to be primarily due to the positive charge characteristics of R9, and not local differences in channel diameter (as described in the previous section).

The K^+^ ion PMF profiles for the designed variants show that replacing the entire NT domain of Cx46 with that of Cx50 (Cx46-50NT) or the 9^th^ residue (Cx46-R9N) substantially reduced/eliminated the differences of NT domain energy barriers between Cx46 and Cx50 (ΔΔG_K+_ = 0 – 0.2 kcal mol^−1^), by effectively reducing the peak barrier around the 9^th^ position (Fig. 3c). Likewise, replacing the entire NT domain of Cx50 with that of Cx46 (Cx50-46NT) or the 9^th^ residue (Cx50-N9R) also reduced the energy difference between Cx46 and Cx50 (ΔΔG_K+_ = 0.2 – 0.5 kcal mol^−1^), however, this time by increasing the peak barrier to a similar level of Cx46 (Fig. 3d). The slightly lower K^+^ energy barrier obtained for Cx50-46NT (ΔG_K+_ = 1.8 kcal mol-1), compared to Cx46 and Cx50-N9R (ΔG_K+_ = 2.1 – 2.3 kcal mol^−1^), is proposed to reflect the slightly larger pore diameter of this model (see Fig. 1d). Taken together, these data confirm that the primary difference in K^+^ energy barriers between Cx46 and Cx50 is due to the placement of a positively charged arginine at the 9^th^ position, and led to the hypothesis that swapping these sites/domains would effectively convert the ion conductance properties of these channels, which was tested by the experiments described in the following sections.

### Cx46-50NT forms non-functional gap junction channels, and is reflected by NT-domain instability observed by MD-simulation

In order to integrate the results of our structural models and MD studies to GJ channel properties, we generated sheep Cx46, Cx50, the NT domain swapped chimeras (Cx46-50NT and Cx50-46NT), and point variants at the 9^th^ residue (Cx46-R9N and Cx50-N9R) and expressed each of them in GJ-deficient N2A cells for functional characterization. N2A cells were transfected with expression vectors containing one of our designed constructs and an untagged GFP (*e.g*., Cx46-IRES-GFP, Cx46-R9N-IRES-GFP, Cx46-50NT-IRES-GFP, Cx50-IRES-GFP, Cx50-N9R-IRES-GFP or Cx50-46NT-IRES-GFP) (Fig. 1a,b). Cell pairs with positive expression of GFP were voltage clamped using dual whole cell patch clamp technique. Approximately half of the cell pairs expressing Cx46 or Cx50 showed junctional currents (I_j_s) in response to a V_j_ pulse (Fig. 4a,b), indicating successful formation of functional GJs. Cell pairs expressing the chimera, Cx50-46NT or one of the point variants (Cx46-R9N and Cx50-N9R) were also frequently coupled with similar coupling percentages as those of wildtype lens connexins (Fig. 4a,b). However, none of the cell pairs expressing Cx46-50NT showed any I_j_s in 6 independent experiments (Fig. 4a,b). The average coupling conductance (G_j_) for each expressed construct was calculated from the GJ coupled cell pairs and no significant differences were observed among different constructs with the exception of Cx46-50NT (Fig. 4c).

**Fig. 4.**
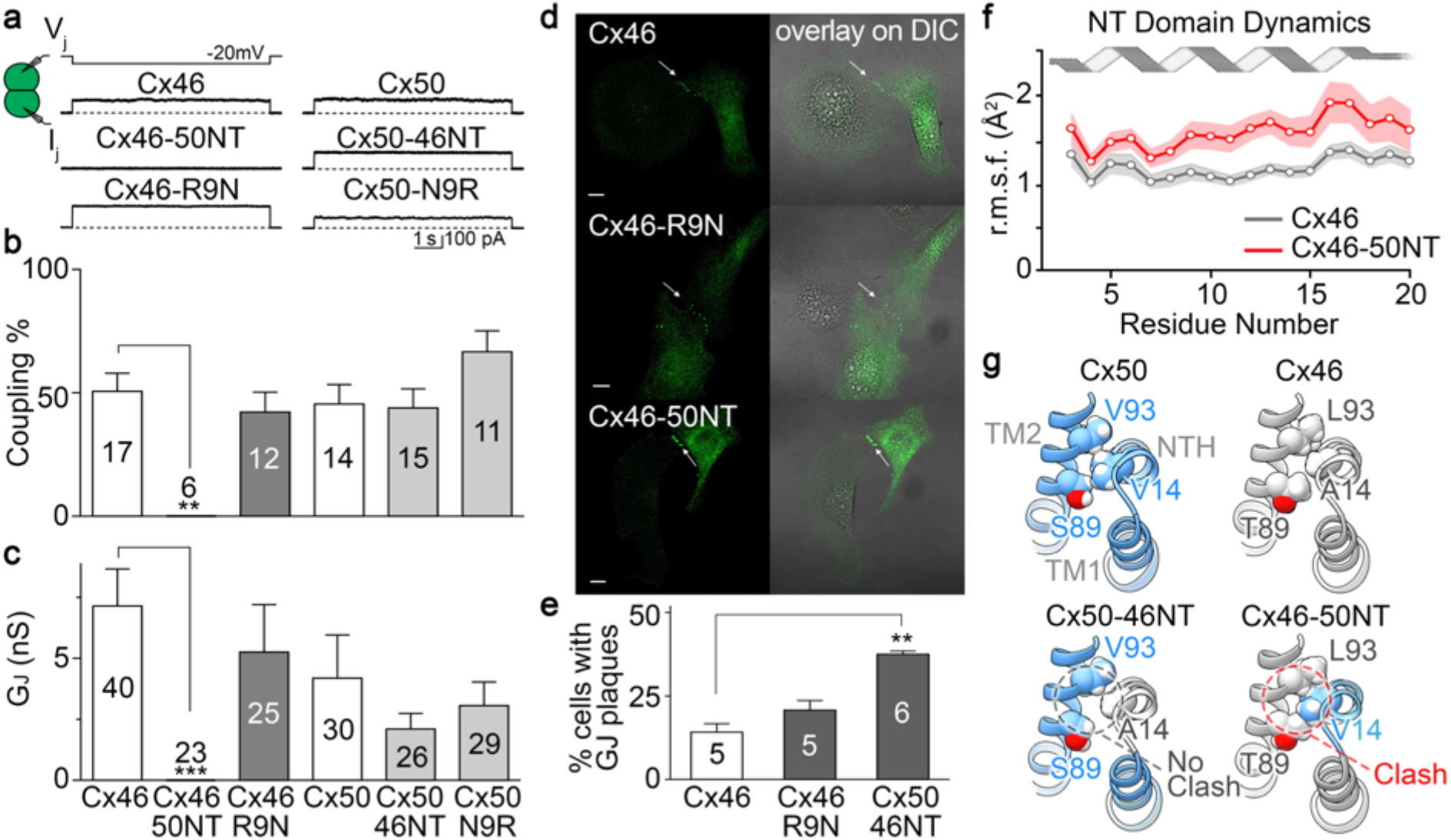
GJ channel function, cellular localization and NT domain stability of Cx50, Cx46 and designed NT domain variants. **a.** Dual whole-cell patch clamp technique was used to measure junctional current (I_j_) in N2A cell pairs expressing GFP-tagged constructs of sheep Cx50, Cx46 and NT domain variants (Cx50-N9R, Cx50-46NT, Cx46-R9N and Cx46-50NT) in response to a −20 mV V_j_ pulse. **b.** Bar graph showing the average coupling percentages of cell pairs expressing homotypic sheep Cx50, Cx46 and designed NT domain variants. One-way ANOVA followed by Newman-Keuls post-hoc test was used compare each of the variants with their respective controls. Cx46-50NT showed no coupling, which was significantly different from that of wildtype Cx46 (** P < 0.01). The number of transfections is indicated. **c.** Bar graph showing the average coupling conductance (G_j_) of coupled cell pairs. Kruskal-Wallis followed by Dunn’s post hoc test was used to compare each of the variants with their respective controls. The total number of cell pairs is indicated. d. Cell imaging of GFP-tagged Cx46, Cx46-R9N and Cx46-50NT expressed in connexin deficient HeLa cells. GJ plaque-like structures (arrows) similar to that of Cx46-GFP (upper panels) are identified for both Cx46-R9N and Cx46-50NT variants. GFP fluorescent signals are superimposed onto DIC images to show the localization of these fluorescent signals (panels on the right). Scale bars = 10 μm. **e.** Bar graph showing the percentage of cells expressing each of the constructs displaying GJ plaque-like structures at the cell-cell interfaces. Note that the Cx46-50NT showed a significantly higher percentage to observe GJ plaque-like structures (**P < 0.01). **f.** Line graph showing NT domain dynamics obtained by MD simulation, as assessed by the backbone root mean square fluctuation (r.m.s.f.) for Cx46 (gray trace) and Cx46-50NT (red trace). Shaded boundaries indicate the 95% confidence intervals. Residues 3 – 20 show significant difference between these models (P < 0.05). **g.** Structural comparison of hydrophobic packing interactions involving residue 14 on the NT domain and residues 89 and 93 on TM2 of Cx50, Cx46 and chimeric models for Cx50-46NT and Cx46-50NT. The Cx46-50NT chimera introduces a steric clash involving bulky residues V14 and T89.

To test if the failure of Cx46-50NT to form functional GJs was due to impairment in the localization of this chimera, we used GFP fusion tagged at the carboxyl terminus of Cx46-50NT. As shown in Fig. 4d,e, Cx46-50NT-GFP was localized in intracellular compartments and displayed GJ plaque-like clusters at the cell-cell interfaces with a higher percentage than that of Cx46-GFP, suggesting that it is unlikely due to abnormal localization of this chimera for its failure in forming functional GJs. For comparison, Cx46-R9N-GFP showed a similar percentage of forming GJ plaques as that of Cx46-GFP (Fig. 4d,e).

To gain insight into the molecular basis for the loss of GJ channel function in the Cx46-50NT chimera, we further interrogated the MD simulation data for this construct. Notably, the dynamical behavior of the NT domain in this construct is significantly higher compared to the wildtype Cx46 model, as assessed by the backbone root-mean-square-fluctuation (r.m.s.f.), which describes the amplitude of backbone dynamics (Fig. 4f). The reason for the decreased conformational stability of the NT domain in this chimera is suspected to be due to the introduced changes in hydrophobic anchoring residues within the NT domain (Fig. 1a,b). Specifically, placement of the Cx50 NT domain onto the Cx46 channel introduces an apparent steric clash between V14 and T89 (located on TM2) (Fig. 4g). In Cx50, position 89 is occupied by a small serine residue, which can accommodate a bulky V14 anchoring residue. In contrast, Cx46 appears to compensate for the smaller A14 anchoring site with a bulkier T89 residue. Of further comparison, when the Cx46 NT domain (containing the smaller A14 site) is placed onto the Cx50 channel (containing the smaller S89 site), a vacant space is introduced between the NT domain and TM2 of the Cx50-46NT chimera (Fig. 4g). This arrangement allows the NT domain to pack more closely against TM2 during MD simulation, resulting in the larger pore diameter of this construct as compared to Cx46, Cx50 and other variant models (see Fig. 1d).

### Swapping the NT-domain or the 9^th^ residue of Cx46 onto Cx50 alters V_j_-gating, while V_j_-gating of Cx46-R9N is unaffected

To investigate the effects of NT domain variants on transjunctional voltage dependent gating (V_j_-gating), cell pairs forming homotypic GJs were recorded by dual whole cell patch clamp and their I_j_s were measured in response to a series of V_j_ pulses from ±20 to ±100 mV (Fig. 5a,b). The I_j_s of Cx46 and Cx50 GJs both showed symmetrical V_j_-dependent deactivation, with similar V_j_-gating properties (Fig. 5a). For both wildtype GJs, I_j_s in response to V_j_s in the range of ±40 to ±100 mV, showed strong deactivation (Fig. 5a). When V_j_ absolute value was ≤ 20 mV, I_j_s showed no deactivation (Fig. 5a). The normalized steady state conductance (G_j,ss_) was plotted as a function of V_j_ (Fig. 5b), which could be well fitted by a Boltzmann equation for each V_j_ polarity, for either Cx46 or Cx50 GJs (Fig. 5b). None of the Boltzmann fitting parameters of Cx46 and Cx50 were significantly different (Table 1).

**Fig. 5.**
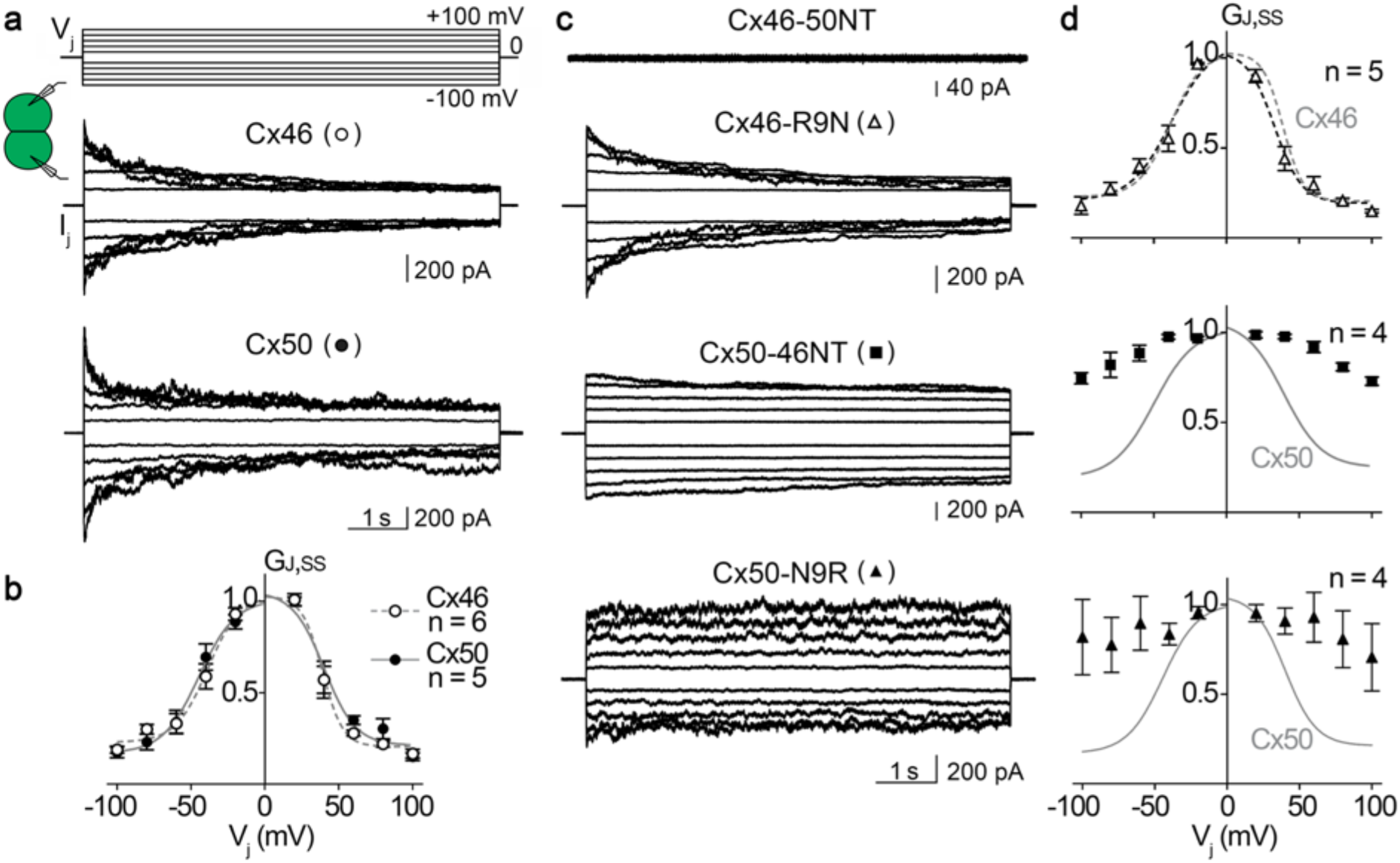
V_j_-gating of homotypic sheep Cx46, Cx50 and NT domain variant gap junction channels. **a, c.** Superimposed junctional currents (I_j_s) recorded from cell pairs expressing homotypic sheep Cx46 or Cx50 GJs (panel a) and NT domain variants Cx46-50NT, Cx46-R9N and Cx50-N9R GJs (panel c), in response to a series of V_j_ pulses (shown on the top of panel a, ±20 to ±100 mV with 20 mV increment). **b.** Normalized steady state junctional conductance, G_j,ss_, of Cx46 (open circles) and Cx50 (filled circles) plotted as a function of V_j_s. Boltzmann equations were used to fit G_j,ss_ – V_j_ plots for Cx46 (smooth dashed grey lines) and Cx50 (smooth solid grey lines) GJs. **d.** Normalized steady state junctional conductance, G_j,ss_, of Cx46-R9N (open triangles), Cx50-46NT (filled squares), and Cx50-N9R (filled triangles) plotted as a function of V_j_s. Only the G_j,ss_ – V_j_ plot of Cx46-R9N GJ was fitted well with Boltzmann equations (smooth dashed black lines). For comparison, the Boltzmann fitting curves of wildtype Cx46 (smooth dashed grey lines) or Cx50 GJ (smooth solid grey lines) were superimposed on the respective plot. The number of cell pairs for each construct is indicated in panels b and d.

**Table 1.** Boltzmann fitting parameters for homotypic sheep Cx46, Cx50, and Cx46-R9N GJs

|  | **V<sub>j</sub> polarity** | **G<sub>min</sub>** | **V<sub>0</sub>** | **A** |
| --- | --- | --- | --- | --- |
| Cx46<br>(n = 6) | + | 0.21 ± 0.02 | 39.0 ± 1.5 | 0.14 ± 0.04 |
|  | - | 0.23 ± 0.03 | 38.6 ± 2.9 | 0.09 ± 0.02 |
| Cx50<br>(n = 5) | + | 0.21 ± 0.04 | 40.1 ± 3.2 | 0.09 ± 0.02 |
|  | - | 0.17 ± 0.04 | 44.5 ± 3.4 | 0.08 ± 0.02 |
| Cx46-R9N<br>(n = 5) | + | 0.19 ± 0.03 | 33.2 ± 2.7 | 0.11 ± 0.03 |
|  | - | 0.21 ± 0.05 | 38.0 ± 4.4 | 0.07 ± 0.02 |

Data is presented as mean ± SEM, and V<sub>0</sub> are absolute values. One-way ANOVA followed by Tukey post-hoc test was used to compare Boltzmann fitting parameters of all three GJs. None of the parameters showed statistically significant differences for the same V<sub>j</sub> polarity.

V_j_-gating of Cx46-50NT, Cx46-50R9N, Cx50-46NT, Cx50-N9R GJs were then studied using the same V_j_ protocol (Fig. 5c,d). As shown in the Fig. 5c, Cx46-50NT was unable to form functional GJs, while Cx46-R9N, Cx50-46NT, and Cx50-N9R all successfully formed functional GJs. Among these functional GJs, Cx46-R9N showed strong symmetrical deactivation when V_j_s were in the range of ±40 to ±100 mV (Fig. 5c). The G_j,ss_ – V_j_ plot for Cx46-R9N could be well fitted with the Boltzmann equation for each V_j_ polarity, which showed no significant difference from those of wildtype Cx46 GJs (Fig. 5d) and Table 1. The I_j_s of Cx50-46NT GJs showed very weak deactivation in the tested V_j_s (Fig. 5c), and the G_j,ss_ – V_j_ plot of these channels could not be fitted with a Boltzmann equation (Fig. 5d). Similarly, I_j_s of Cx50-N9R GJ also showed no consistent deactivation in the tested V_j_s (Fig. 5c), and the G_j,ss_ – V_j_ plot also could not be fitted with a Boltzmann equation for these channels (Fig. 5d). In addition, the I_j_s of Cx50-N9R showed a lot of fluctuations throughout the V_j_ pulse (Fig. 5c) resulting in large variations in the G_j,ss_ (Fig. 5d). Taken together, these data indicate that the V_j_-response for variants of Cx50 (Cx50-46NT and Cx50-N9R) are more susceptible to perturbation than the Cx46 variant (Cx46-R9N). A structural basis for these differences was not entirely clear, as the millisecond-second long time-scales of V_j_-gating are beyond the limits of traditional MD simulation (see Discussion).

### Differences in Cx50 and Cx46 GJ single-channel conductance properties are defined by the 9^th^ residue

MD simulation studies indicated that the incorporation of the positively charged arginine at the 9^th^ position of Cx46, Cx50-N9R and Cx50-46NT introduces a significant energetic barrier to K^+^ ion permeation (the major permeant ion of Cx46/Cx50 GJs), as compared to Cx50 and Cx46-R9N (see Fig. 3). To investigate the functional effects of this positively charged residue we characterized the unitary conductance (γ_j_) of Cx50, Cx46 and the designed NT domain variants based on single channel current (i_j_) recordings at varying V_j_s. In Fig. 6a, homotypic sheep Cx46 and Cx50 single channel current (i_j_) traces are shown at the V_j_s indicated. All point histograms and Gaussian fits were used to measure the amplitudes of i_j_s for the main open state at the tested V_j_s (Fig. 6b). The averaged i_j_s were plotted at different V_j_s and a linear regression i_j_ – V_j_ plot for Cx46 or Cx50 GJs was used to estimate the slope unitary channel conductance (γ_j_) (Fig. 6c). The γ_j_ of Cx46 GJ is 170 ± 3 pS, whereas the γ_j_ of Cx50 GJ is 208 ± 3 pS.

**Fig. 6.**
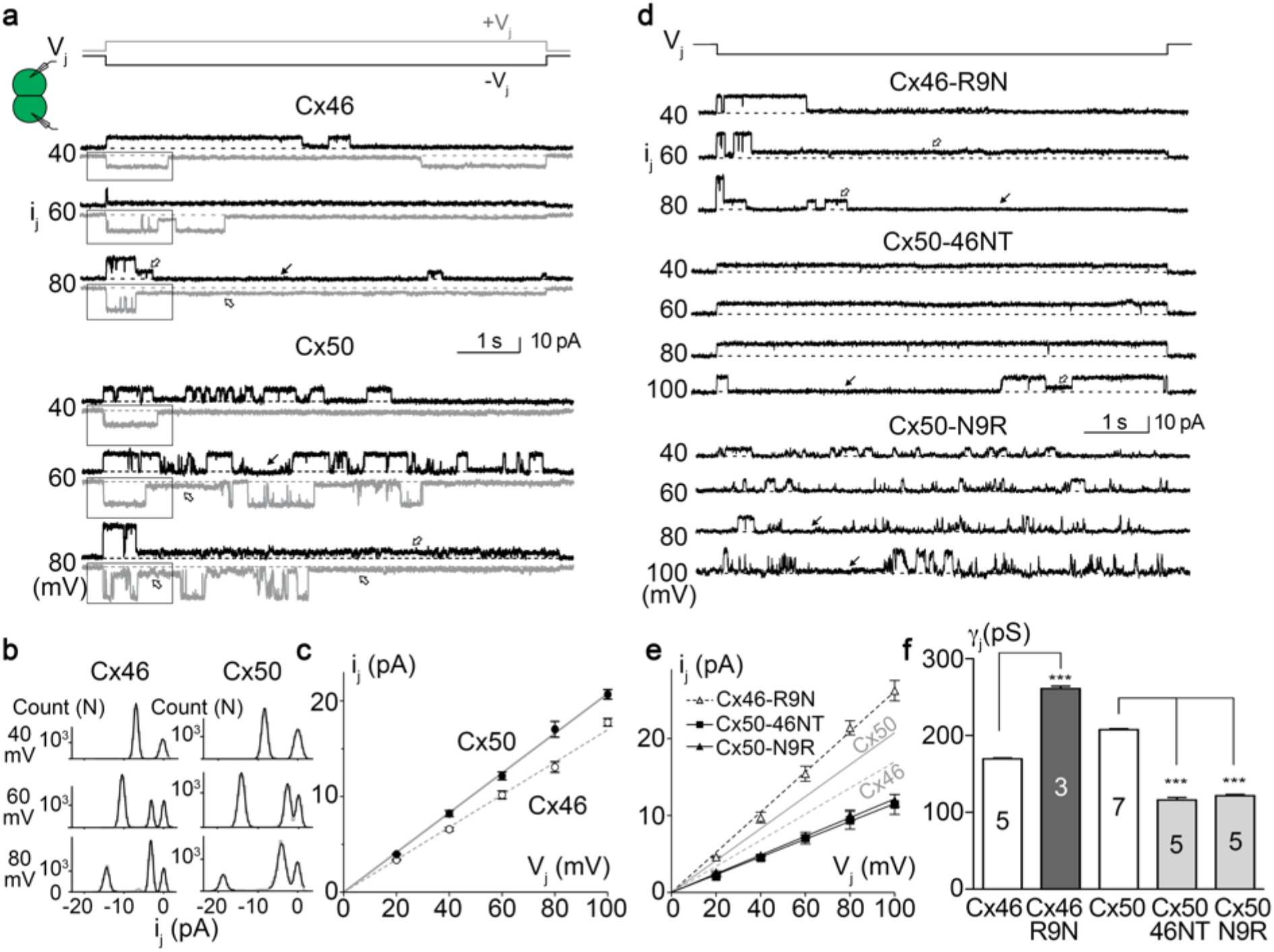
Single channel properties of homotypic sheep Cx46, Cx50 and NT-domain variant GJs. **a.** Single channel currents (i_j_s) recorded from a cell pair expressing wildtype sheep Cx46 (top set) or Cx50 (bottom set) at the indicated V_j_s. Positive V_j_ (+V_j_) or negative V_j_ (-V_j_) induced i_j_s are shown in grey or black, respectively. Both GJs showed fully open state at the beginning of V_j_ pulses and transitions to either subconductance (open arrows) or fully closed state (solid arrows). Note that more than one subconductance states could be identified at ±80 mV V_j_s. **b.** All point histogram and Gaussian fits were used to estimate the i_j_ amplitude at the main open state. A selected section as shown in panel a was used to generate all point histogram. Main conductance state and in some cases a subconductance state could be identified at different V_j_s. **c.** The i_j_ amplitudes of the main open state were plotted at each tested V_j_. The slope of the linear regression lines in the i_j_ – V_j_ plot represent the slope unitary channel conductance (γ_j_). For Cx46 GJ, the slope γ_j_ = 170 ± 3 pS (n = 5) and for Cx50, the slope γ_j_ = 208 ± 3 pS (n = 7). **d.** Single channel currents (i_j_s) recorded from cell pairs expressing sheep Cx46-R9N, Cx50-N9R, or Cx50-46NT at the indicated V_j_s. The i_j_s of Cx46-R9N showed a main open state at the beginning of the V_j_ pulses and transitioned into subconductance state (open arrows) or fully closed state (solid arrow). The i_j_s of Cx50-46NT preferentially reside at fully open state at most tested V_j_s (−40 to −80 mV) and transition to fully closed state (solid arrow) or to a subconductance state could be observed at −100 mV V_j_. The i_j_s of Cx50-N9R showed multiple open events with frequent transitions to fully closed state (solid arrows). **e.** The amplitude of i_j_s for the main open state at each tested V_j_ was plotted against V_j_ and analyzed as described for panel c. The solid and dashed grey lines represent the slope γ_j_s of Cx50 and Cx46 respectively. The slope γ_j_ of Cx46-R9N is 261 ± 6 pS (n = 3), Cx50-N9R γ_j_ is 122 ± 3 pS (n = 5), and Cx50-46NT γ_j_ is 116 ± 5 pS (n = 5). f. Bar graph to summarize the slope γ_j_s of all tested variants. The γ_j_ of Cx46-R9N was significantly higher than that of Cx46 and the γ_j_s of Cx50-46NT and Cx50-N9R were significantly lower than that of Cx50 (one-way ANOVA followed by Tukey’s post hoc test for biological meaningful groups).

Fig. 6d shows i_j_s of homotypic Cx46-R9N, Cx50-N9R, and Cx50-46NT GJs at the indicated V_j_s. The averaged i_j_s for the main open state for each tested V_j_ of these variants were plotted to create an i_j_ – V_j_ plot. Linear regression of i_j_ – V_j_ plot for each variant was used to estimate slope γ_j_ (Fig. 6e). Cx46-R9N GJs showed a significant increase in slope γ_j_ (261 ± 6 pS) from that of wildtype Cx46 GJ (Fig. 6e). Conversely, Cx50-N9R GJs displayed a significantly decreased slope γ_j_ (122 ± 3 pS) than that of wildtype Cx50 GJ (Fig. 6e). Replacing the NT domain of Cx50 with that of Cx46, Cx50-46NT, resulted in a more pronounced drop in slope γ_j_ (116 ± 5 pS). The slope single channel conductance of all tested variants were plotted as a bar graph for direct comparison (Fig. 6f). These effects are qualitatively consistent with the predicted results obtained by MD, however there was a surprisingly higher degree of effect of the placement/removal of R9 leading to channel variants with correspondingly higher or lower unitary conductance, as compared to wildtype channels.

### Cx50-46NT forms GJ channels with an extremely stable open-state, while Cx50-N9R open-state becomes destabilized

Another surprising result revealed by single channel recordings was the effects on the open state probability of the variant GJs. As shown in Fig. 6d, the i_j_s of Cx50-46NT were virtually in the fully open state (or probability of open, P_open_ ≈ 1) at all V_j_s except at 100 mV, whereas i_j_s of Cx50-N9R show a much lower P_open_ at all tested V_j_s (P_open_ in the range of 0.11 - 0.22). The i_j_s of both Cx50-46NT and Cx50-N9R showed little change in P_open_ in the tested V_j_s (data not shown).

Upon close inspection of the i_j_s of Cx46, Cx50 and their variants, we found that the open dwell times were different among different GJs. To quantify open dwell time for each of these GJs, the i_j_s of these GJs were further analyzed to obtain open dwell time for each open event (Fig. 7a). At V_j_ of −60 mV, a limited number of open events (transitions from closed to open and return to the closed state) could be identified for Cx50-46NT during a 7-second V_j_ pulse. A few open events for Cx46 and Cx46-R9N GJs and many more open events for Cx50 and Cx50-N9R GJs were detected at the same V_j_ (Fig. 7a). The open dwell time for all events was averaged for each variant GJ (Fig. 7b). In most of the tested V_j_s, the rank of the open dwell time from long to short is in the following order: Cx50-46NT > Cx46 > Cx50 > Cx46-R9N > Cx50-N9R GJs (Fig. 7b).

**Fig. 7.**
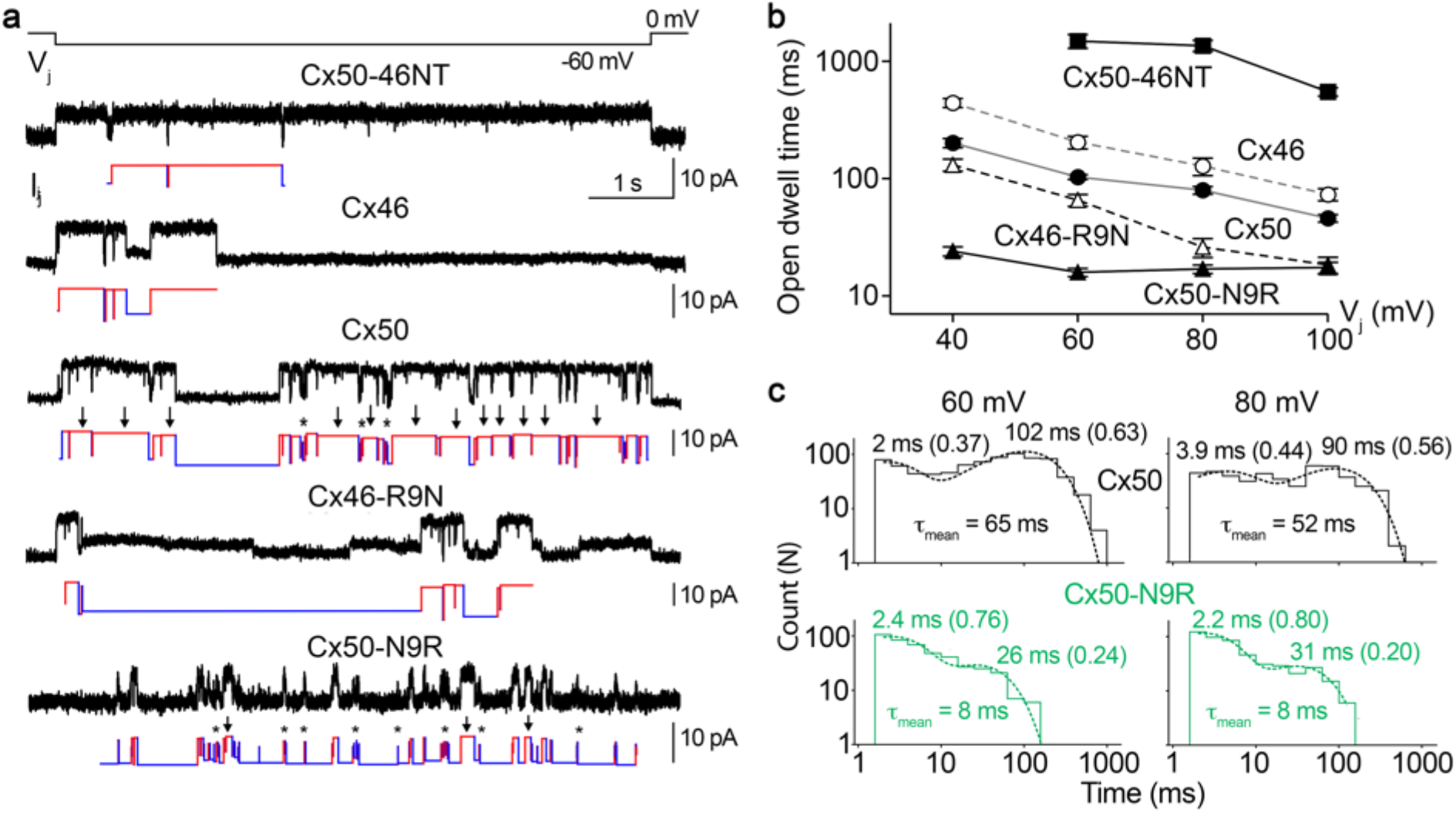
Single channel open state dwell times of Cx50, Cx46 and NT domain variants. **a.** The i_j_s (black) and schematic single channel openings (red) and closings (blue) are shown for sheep Cx50-46NT, Cx46, Cx50, Cx46-R9N, Cx50-N9R GJs. Cx50 and Cx50-N9R GJs showed apparently short (asterisks, *) and long (arrows) open events, as well as more open events than the other GJs. Note that the first and the last open events were excluded as they likely under represent the open dwell time (*e.g*. exist in open state before or after the V_j_ pulse). **b.** The averaged open dwell time for each GJ were plotted at different V_j_s. The number of events analyzed ranged from 33-832. Note that the open dwell time is plotted in logarithmic scale to show large differences among these GJs. The ranking of the open dwell times of these GJs from high to low is: Cx50-46NT > Cx46 > Cx50 > Cx46-R9N > Cx50-N9R. **c.** The open dwell time for all open events for Cx50 and Cx50-N9R were plotted in a logarithmic histogram with 5 bins/decade and fitted with two-exponential log components. The two time constant (τ) values represent the corresponding short and long open events for Cx50 (top histograms) and Cx50-N9R (bottom green histograms). The τ_mean_ values were calculated by taking the weighted average of the two τ values using the area under each peak for each of these variants at a specific V_j_.

It is interesting to note that open event frequency was much higher for both Cx50 and Cx50-N9R GJs and both GJs showed two types of openings, short (labeled by asterisks) and long (labeled by arrows) opening events (Fig. 7a). Histograms of open dwell time were generated for both of these GJs (Fig. 7c) and each of them could be fitted with two-exponential components. Cx50 showed short (2 ms) and long (102 ms) open events at V_j_ 60 mV with a higher prevalence of the long open events (63%) (Fig. 7c). At V_j_ of 80 mV, the average of long open events became shorter (90 ms) with slightly lower percentage (56%), while the average of the short open events is 3.9 ms with an increased percentage (44%). Weighted average of time constants (τ_mean_) for these two V_j_s were also calculated (Fig. 7c). Upon mutating the 9^th^ residue to arginine (Cx50-N9R) the long open events became much shorter (26 ms and 31 ms) with reduced percentages (24% and 20%), respectively for V_j_s of 60 and 80 mV. The Cx50-N9R τ_mean_ value was much lower than that of Cx50. Taken together, these results suggest that the identity of the 9^th^ residues, in addition to other structural differences within the NT domains of these variants significantly influence the open-state probability of these GJs (see Discussion).

## Discussion

In this study, structural modeling and molecular dynamics simulations were conducted in combination with macroscopic and single channel electrophysiology studies, to investigate the mechanistic roles of NT domain in two closely related lens GJs, Cx50 and Cx46. We identified differences in structure and dynamics within the NT domains, in particular the 9^th^ position, as the key features differentiating the pore profiles, energetic barriers to ion permeation, ion conductance properties and open-state stabilities between these two isoforms (see summary of results presented in Fig. 8). The CryoEM-based models of Cx46 and Cx50 indicate that the NT domains of these connexins formed constriction sites at the entrances of a GJ channel (Myers *et al*., 2018) and displayed electrostatic properties that could directly influence the efficiency of substrate permeation (*e.g*., due to differences in size, shape, and charge properties etc.) and the rate of ion permeation, which is directly measurable as γ_j_ using dual patch clamp. All-atom equilibrium MD simulation on Cx46, Cx50, Cx50-46NT, Cx46-R9N, and Cx50-N9R GJs suggested that all of these channels displayed much higher free energy barrier (described by PMF) for Cl^-^ (the major intracellular anion) versus K^+^ (the major intracellular cation), making these GJs more permeable to cations with a difference in free energy barriers ΔΔG_(Cl- – K+)_ = ~2.4 – 3.7 kcal mol^−1^. These results are consistent with previous electrophysiology studies conducted on rodent Cx50 and Cx46 that demonstrate K^+^/Cl^-^ permeability ratios (*P*_K+_/*P*_Cl-_) of ~7/1 (Trexler *et al*., 1996; Srinivas *et al*., 1999; Trexler *et al*., 2000; Tong *et al*., 2014). At the same time, the differences in the peak free energy barrier to K^+^ permeation between Cx50 and Cx46 (ΔΔG_K+_ = 0.8 kcal mol^−1^) is consistent with the relative differences in single channel conductance for these isoforms, where we showed sheep Cx50 displays higher conductance compared to Cx46 (208 pS vs 170 pS).

**Fig. 8.**
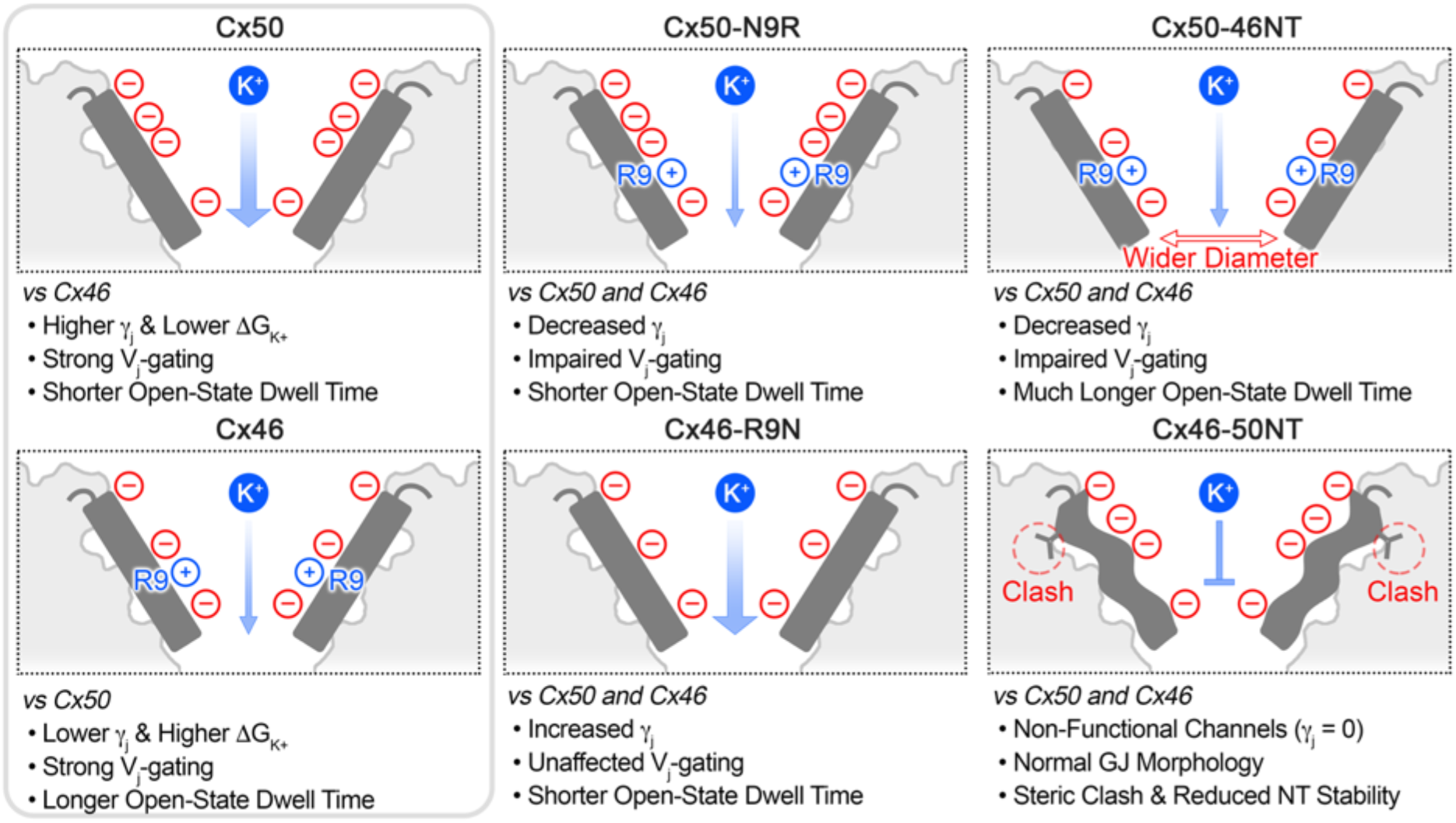
Overview of significant structural and functional differences between Cx50, Cx46 and NT domain variants. Cartoon illustrating the NT domain regions (shown in dark grey) and structural and/or functional differences between Cx50, Cx46 and the designed variants for this study. Locations of negatively charged residues (red circles) and positively charged R9 residue (blue circles) are indicated (see for comparison Fig. 1b). The relative permeabilities to the major permeant ion (K^+^) is indicated by the width of the blue arrow, where Cx50 displays a higher unitary conductance and lower ΔG_K+_, as compared to Cx46. Both wildtype GJs displayed strong V_j_-gating. The single point variants, Cx50-N9R and Cx46-R9N, produced channels with γ_j_s that were augmented by adding or removing the positively charged R9 position, although these variants affected V_j_-gating and open state dwell times differently. The Cx50-46NT chimera displayed remarkably long open-state dwell times and impaired V_j_ sensitivity, as compared to wildtype GJs, which correlated with a tighter packing of the NT domain to TM2 resulting in a wider pore radius compared to Cx46 or Cx50 following MD simulation. The Cx46-50NT construct formed morphological GJs, but these channels were non-functional. The loss of channel activity for this construct is proposed to be due to a steric clash that is introduced between V14 and T89 and reduced stability of the NT domain observed by MD simulation (indicated by the wavy NT domain in the illustration).

The differences in peak energy barrier to K^+^ and single channel conductance observed among Cx50, Cx46 and the NT variants appear to be most directly linked to the charge property and orientation of their 9^th^ position (see Fig. 8). In the CryoEM-based model of Cx46, the positively charged R9 residue is oriented toward the center of the pore and forms a narrow constriction site that is absent in Cx50 (see Fig. 1a,c and 2a). However, it is noted that the conformation of this residue is not well defined by the CryoEM density map (Myers *et al*., 2018). Indeed, during MD simulation, R9 shows significant fluctuations and appears to stably adopt multiple conformational states that reorient this bulky sidechain away from the pore and positioned against the lumen of the channel through inter-subunit interactions with E12 and/or a neighboring R9 site (see Fig. 2b-d). The effective result of this behavior is Cx50, Cx46 and each of the designed NT domain variants (except Cx50-46NT) all displayed very similar pore profiles during MD simulation (Fig. 1d), with a minimum diameter of ~9.2 – 9.6 Å localized around S5/D3. Cx50-46NT displayed a slightly wider pore diameter of ~11 Å (discussed below). These dimensions are consistent with previous experimental permeability studies that indicate GJ channels are permeable to substrates of ~8 – 11 Å diameter (Trexler *et al*., 1996; Gong & Nicholson, 2001; Kanaporis *et al*., 2008), and is sufficient to support the permeation of a wide-variety of hydrated ions and small molecule substrates.

The free energy landscape to cation permeation was similar between GJ variants, depending on the presence or absence of a positively charged arginine at the 9^th^ position. For example, the peak ΔG_K+_ for the variants Cx46-R9N and Cx46-50NT were both reduced to that of Cx50, and the peak ΔG_K+_ for Cx50-N9R and Cx50-46NT variants were increased to a degree that was similar to Cx46 (see Fig. 3). By functional comparison, our dual patch clamp data on these variants demonstrate that the γ_j_s of these variants displayed unitary conductances in order from high to low: Cx46-R9N (263 pS) > Cx50 (208 pS) > Cx46 (170 pS) > Cx50-N9R (122 pS) = Cx50-46NT (116 pS), while Cx46-50NT was non-functional (discussed below). These results are consistent with free energy calculations obtained by MD, at least on a qualitative level. Surprisingly, however, Cx46-R9N γ_j_ was much higher than that of Cx50 and the γ_j_s of Cx50-N9R and Cx50-46NT were much lower than that of Cx46. The mechanistic basis for this observation is not entirely clear, but could be due to one or combinations of the following factors: 1) contributions to channel conductance not captured by our structural models (*e.g*., CL and CT domains were not modeled due to intrinsic disorder of these domains); 2) cation/anion permeation ratio could be altered by the variants; 3) inherent limitations in the force-fields and/or time-scales used for MD simulation. Future experiments are needed to resolve some of these possibilities.

The chimeras with the NT domain (the first ~20 residues) switched between Cx46 and Cx50 were either unable to form functional GJ channels (Cx46-50NT) or formed functional channels (Cx50-46NT) with altered V_j_-gating, reduced γ_j_, and increased channel open dwell time (a parameter for stability of the open-state). Remarkably, Cx46-50NT GJs appeared to be able to reach the cell membrane and formed morphological GJs (see Fig. 4d,e), indicating that the loss of channel function was unlikely due to defects in trafficking or GJ assembly. Possible mechanistic insights to this phenotype are provided by our structural models, where this chimera introduces a specific steric clash between the introduced V14 on the NT domain and T89 located on TM2 (Fig. 8 and see Fig. 4g). During MD simulation, Cx46-50NT displayed enhanced dynamical behavior within the NT domain, exemplified by r.m.s.f. amplitudes that were significantly larger than Cx46 (see Fig. 4f). Taken together, these data suggest that the NT domain of Cx46-50NT becomes destabilized, likely by steric interactions involving hydrophobic anchoring sites, resulting in loss of channel function (Fig. 8). Although, improper folding of the Cx50 NT domain in the context of this chimera can also not be ruled out. In contrast, placement of the Cx46 NT domain onto Cx50 resulted in more close packing (due to smaller hydrophobic residues at positions 14 and 89), which resulted in an overall wider pore diameter for this construct, as compared to the other GJs under investigation. We suggest that this close packing results in a more stabilized NT domain and contributes, at least in part, to the increased open dwell time and altered V_j_-gating of this GJ. It is interesting to note that a previous study on switching NT domains of mouse Cx50 and rat Cx46 also showed a decrease in hemichannel unitary conductance in Cx50-46NT and surprisingly the hemichannel of Cx46-50NT was functional (unlike our result that sCx46-50NT was unable to form functional GJs) with a slightly higher hemichannel unitary conductance (Kronengold *et al*., 2012). The basis for this difference is not apparently clear, but may reflect the differences in species and/or in the stability of the NT domain in GJ versus hemichannel assemblies.

Both Cx46 and Cx50 GJs displayed prominent V_j_-gating in the range of our tested V_j_s (±20 to ±100 mV) that could be well described by a two state Boltzmann equation for each V_j_ polarity. A classic voltage-gating model for voltage-gated ion channels is that during the gating process, a change in the voltage results in movement of charge or a re-orientation of dipoles (the voltage sensor) relative to the electric field, which leads to either closing or opening of the channel (Bezanilla, 2000). In the case of GJ channels, the sensor for V_j_-gating is believed to reside in the pore lining residues (Harris *et al*., 1981; Bukauskas & Verselis, 2004; Bargiello & Brink, 2009). Removing a positively charged residue in the pore lining NT domain of Cx46 by the variant R9N, showed no change in the V_j_-gating sensitivity (*A*, a Boltzmann parameter can be converted into gating charge, z) and in the half deactivation V_j_ (V_0_), indicating that 1) the positive charge on R9 is unlikely to play a role in V_j_-sensing or V_j_-gating, and 2) there is no change in the free energy difference (ΔG_0_ = zFV_0_) between aggregated open and closed states in the absence of a voltage field (Yifrach & MacKinnon, 2002; Sukhareva *et al*., 2003; Xin *et al*., 2012a). The most striking change associated with Cx46-R9N GJ was an over 50% increase in the γ_j_.

In contrast to the findings on Cx46-R9N, both Cx50-46NT and Cx50-N9R GJs nearly completely lost their V_j_-gating with very little V_j_-dependent deactivation on macroscopic junctional currents (see Fig. 5). We believe that the underlying mechanisms of their apparent loss of V_j_-gating are different based on their single channel gating properties. In the case of Cx50-46NT GJ, the single channel currents (i_j_s) showed long stable opening for all our tested V_j_s with very brief transitions to a residual or closed state. Open probability of Cx50-46NT was approximately one, with longer dwell time in the closed or residual states not observed unless at the highest V_j_s (±100 mV). Quantitative measurements of open dwell time of the Cx50-46NT GJ showed they were about 5 – 15 times longer than those of Cx46 and Cx50 GJs (see Fig. 7). These experimental observations are consistent with our proposed model that switching the NT of Cx46 into Cx50 (Cx50-46NT) stabilize open state of the GJ channel. However, in the case of Cx50-N9R GJs, the opposite effects on i_j_ were observed with reduced open stability (a much shorter open dwell time than wildtype Cx50) and low P_open_ levels for all tested V_j_s without any V_j_-dependence. In addition, long lived residue states were lost in Cx50-N9R GJ and during the entire V_j_ pulses, the i_j_s displayed frequent transitions from closed state to two open states (see Fig. 7). These data support a model where the Cx50-N9R GJs in our experimental conditions are already gated (or not recovered from the V_j_-dependent deactivation during the V_j_ pulse interval) characterized with a fully closed state and frequent transitions to short-lived open states. Consistent with this model, we observed significant gating of Cx50-N9R GJ during their first large V_j_ exposure, not the subsequent V_j_s (Yue and Bai unpublished observations). Further experiments are needed to test these models fully.

To our knowledge, this is the first functional characterization of sheep Cx46 and Cx50 GJs with dual patch clamp on macroscopic and single channel currents. Sheep connexins were selected for this study so that we could directly align functional data with the recently described high resolution GJ structure models (Myers *et al*., 2018; Flores *et al*., 2020). From a structural point of view, the sheep orthologs of Cx46 and Cx50 show significant sequence identity in primary amino acid sequence to human (85% for Cx50 and 66% for Cx46), and over the structured domains the sequence identities are even higher (96% for Cx50 and 95% for Cx46). The entire NT domains of Cx46 or Cx50 are identical among sheep, mouse and human (Myers *et al*., 2018). Congruent to these structural similarities, our functional characterizations of sheep Cx46 and Cx50 GJs also revealed channel properties that are similar to those previously reported for rodent or human homolog connexins. First, the V_j_-gating properties are qualitatively similar to those obtained from rat or human Cx46 GJs and mouse or human Cx50 GJs (White *et al*., 1994a; Srinivas *et al*., 1999; Hopperstad *et al*., 2000; Pal *et al*., 2000; Tong *et al*., 2004; Xin *et al*., 2010; Kronengold *et al*., 2012; Abrams *et al*., 2018). Second, at the single channel level, the γ_j_ of Cx50 (208 ± 3 pS) determined in this study is the same as those determined for mouse or human Cx50 GJs (200 – 220 pS) (Srinivas *et al*., 1999; Xin *et al*., 2010; Rubinos *et al*., 2014). In contrast to Cx50, the reports of γ_j_s of Cx46 are more varied, ranging from 128 – 213 pS depending on species or experimental condition (Hopperstad *et al*., 2000; Trexler *et al*., 2000; Sakai *et al*., 2003; Rubinos *et al*., 2014; Abrams *et al*., 2018). The γ_j_ of sheep Cx46 from the present study was 170 ± 3 pS using CsCl-based ICS, which in the similar range as those reported of rat or human Cx46. Third, the single channel open dwell time of Cx50 showed two open states qualitatively similar to those observed for mouse Cx50 GJ (Xin *et al*., 2012b; Tong *et al*., 2014).

In summary, we believe these studies establish the Cx46/50 structural models obtained by CryoEM as archetypes for structure-function studies targeted at elucidating GJ channel mechanism and the molecular basis of disease-causing variants. Our integrative structural and functional approach was able to provide detailed mechanistic insights on how Cx46 and Cx50 GJs control ion permeation, gating, open state stability, and demonstrate how nuanced differences in the NT domains of Cx50 and Cx46 lead to significantly different channel properties. Understanding these differences at a mechanistic level are important to developing our understanding of how various connexins have adapted to their unique physiological roles throughout the body, and how aberrancies introduced by mutation lead to disease. Indeed, the NT domains of Cx46 and Cx50 are hot spots of genetic variation associated with congenital cataracts (Beyer *et al*., 2013; Mackay *et al*., 2014; Micheal *et al*., 2018; Zhang *et al*., 2018; Ye *et al*., 2019). A similar integrative approach will be key to providing mechanistic insights into the etiology of these and other cataract-associated variants and possibly other connexin-linked diseases. Lastly, we note that during the completion of this manuscript, we reported refined structural models of sheep Cx46 and Cx50 that were resolved to 1.9 Å resolution using CryoEM, in a near-native lipid environment (Flores *et al*., 2020). These refined models provide an unprecedented level of detail for connexin family of intercellular communication channels, and are expected to allow even more detailed investigations targeted at understanding the mechanistic principles underpinning the functional differences of these and other closely related GJ channels.

## Acknowledgments

This work was supported by Natural Sciences and Engineering Research Council of Canada (288241 to D.B.), the National Institutes of Health (R35-GM124779 to S.L.R.) (R01-GM115805 to D.M.Z.) and the National Science Foundation (MCB 1715823 to D.M.Z.).

## Author contributions

All authors contributed to the preparation of the manuscript. B.Y. and B.G.H. contributed equally. B.Y. designed, performed and analyzed patch clamp experiments. H.C. designed and generated all variants and designed and performed GFP-tagged Cx46 variant localization experiments. Z.Z. and M.A. participated in patch clamp experiments, localization experiments, and analyzed relevant data. B.G.H. and U.K. developed homology structure models, designed and conducted MD simulations and performed analysis of MD data. D.M.Z. and S.L.R. supervised the MD studies. D.B supervised the electrophysiology and localization studies. S.L.R. and D.B. provided overall oversight to the design and execution of the work.

## Abbreviations

GJ: gap junction
G_j_: gap junctional coupling conductance
γ_j_: single gap junction channel conductance
I_j_: macroscopic transjunctional current
i_j_: single gap junction channel current
Cx46: sheep connexin 46
Cx50: sheep connexin 50
V_j_: transjunctional voltage
MD: molecular dynamics
r.m.s.d.: root mean square deviation
r.m.s.f.: root mean square fluctuation
PMF: potential of mean force

## Supplemental Data: Tables, Figures and Legends

**Supplemental Table 1.** Summary of molecular dynamics (MD) simulation setup and conditions. Each simulation was setup similarly, using an explicit solvent model containing 150 mM KCl in the cytoplasmic space and 150 mM NaCl in the extracellular space and POPC as the lipid for building the two membranes to mimic GJ environment. The experimental models of sheep Cx50 (PDB: 6MHY) and Cx46 (PDB: 6MHQ) were used as starting models (Myers *et al*., 2018), and modified to include NT acetylation at G2 to match expected co-translational modification found in cells (Shearer *et al*., 2008; Wang & Schey, 2009; Myers *et al*., 2018). NT variants and chimera models were derived from the wildtype experimental models using the *mutator* plugin in VMD (Humphrey *et al*., 1996). Following initial minimization steps, all systems were equilibrated for 30 ns at 37° C (310 K), and multiple replicates (N=4) of production (25 ns each) were acquired for analysis.

|  | Cx50 | Cx50-N9R | Cx50-46NT | Cx46 | Cx46-R9N | Cx46-50NT |
| --- | --- | --- | --- | --- | --- | --- |
| **Total Atoms** | 363,722 | 362,542 | 358,692 | 355,524 | 360,201 | 358,570 |
| Solvent | 194,928 | 193,644 | 190,128 | 191,310 | 191,271 | 189,984 |
| Lipids | 133,866 | 133,866 | 133,598 | 129,042 | 133,866 | 133,464 |
| Protein | 34,488 | 34,608 | 34,560 | 34,764 | 34,644 | 34,692 |
| Ions | 440 | 424 | 406 | 408 | 420 | 430 |
| Modelled Residues | 2–97; 154–234 | 2–97; 154–234 | 2–97; 154–234 | 2–97; 142–222 | 2–97; 142–222 | 2–97; 142–222 |
| Modification to model | n-term acetylation | n-term acetylation | n-term acetylation | n-term acetylation | n-term acetylation | n-term acetylation |
| **Simulation Conditions** |  |  |  |  |  |  |
| Simulation Box (Å) | 146 x 146 x 176 | 146 x 146 x 176 | 146 x 146 x 176 | 146 x 146 x 176 | 146 x 146 x 176 | 146 x 146 x 176 |
| Pressure (atm) | 1 | 1 | 1 | 1 | 1 | 1 |
| Temperature (K) | 310 | 310 | 310 | 310 | 310 | 310 |
| Time Step (fs) | 2 | 2 | 2 | 2 | 2 | 2 |
| Equilibration Time (ns) | 30 | 30 | 30 | 30 | 30 | 30 |
| Production Time (ns) | 25 | 25 | 25 | 25 | 25 | 25 |
| Production Replicas | 4 | 4 | 4 | 4 | 4 | 4 |
| Total Production Time (ns) | 100 | 100 | 100 | 100 | 100 | 100 |

**Supplemental Fig. 1.**
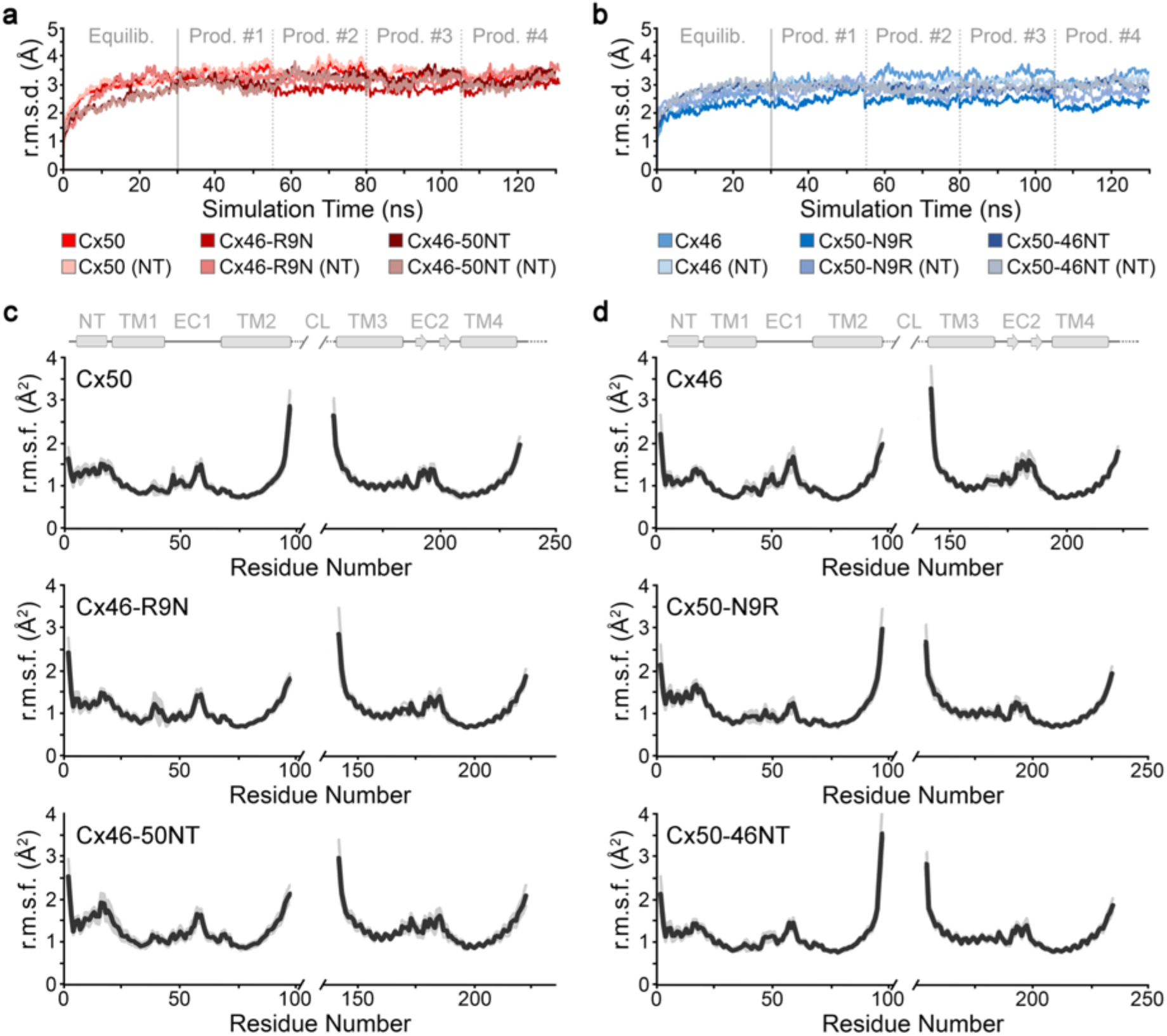
Molecular dynamics setup and validation. **a,b.** Backbone root mean squared deviation (r.m.s.d.) analysis of equilibrium (0 – 30 ns) and production phases (30–130 ns) of the MD simulations, calculated with respect to the experimental starting structures, where Cx50, Cx46-R9N and Cx46-50NT (red traces) are shown in panel a, and Cx46, Cx50-N9R and Cx50-46NT (blue traces) are shown in panel b. Separate analysis for the n-terminal (NT) domains are shown in lighter shades. **c,d.** Plot of average backbone root mean squared fluctuations (r.m.s.f.) displayed during the production phase of the molecular dynamics (MD) simulations for Cx50, Cx46-R9N and Cx46-50NT (panel c) and Cx46, Cx50-N9R and Cx50-46NT (panel d). Averages are determined for the 12 subunits composing the intercellular channel, analyzed for the four independent productions. Error bars (light grey shading) represent 95% confidence intervals (N = 12). Secondary structure and domain labels are indicated for the NT domain, transmembrane helices (TM1-4), extracellular domains (EC1-2) and cytoplasmic loop (CL; not modeled).

**Supplemental Fig. 2.**
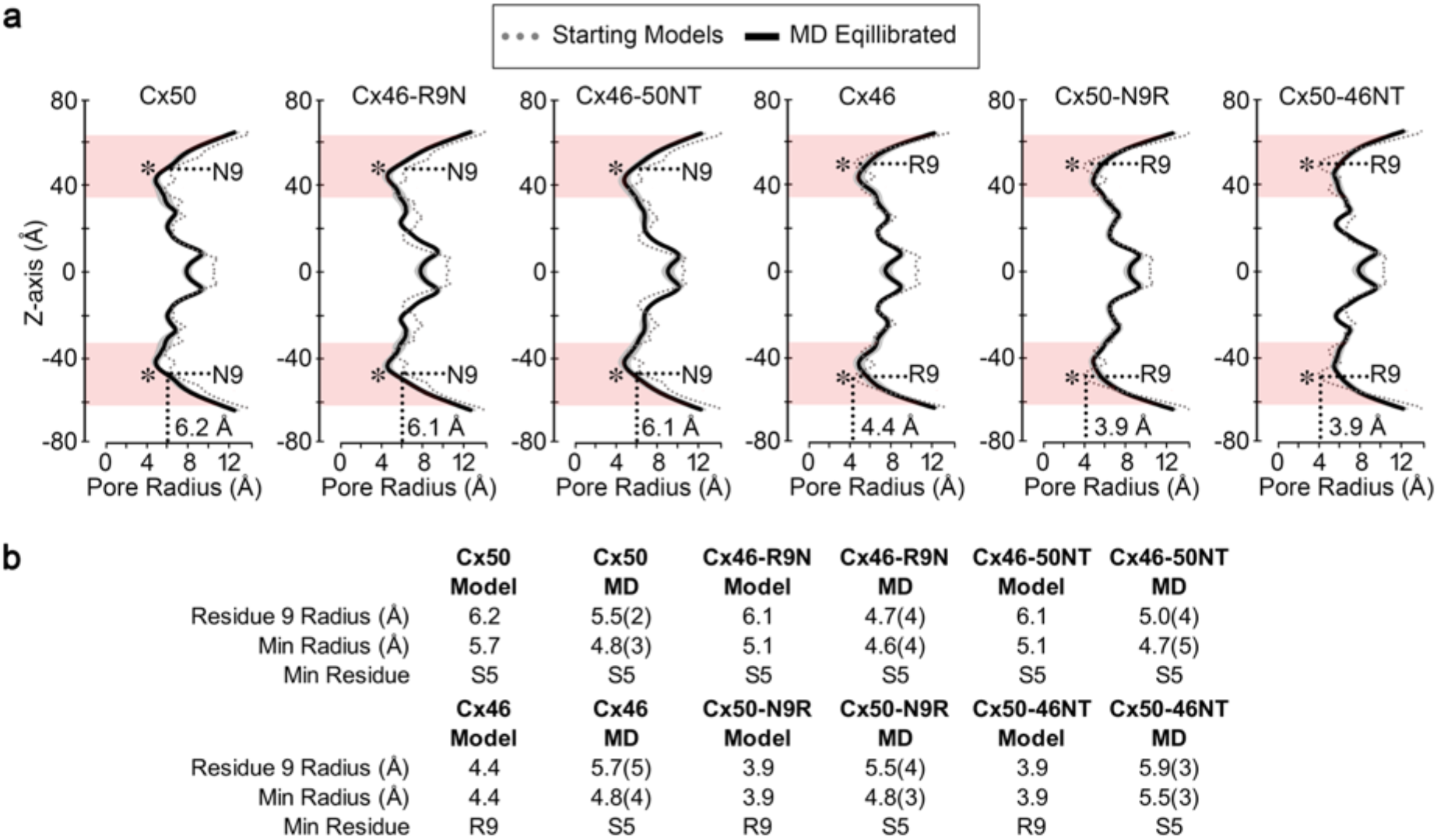
Pore profile analysis of Cx50, Cx46 and designed variants. **a.** Full representation of pore radius diagrams of Cx50, Cx46 and designed variants obtained using the program HOLE (Smart *et al*., 1996). Pore profiles corresponding to the experimental starting structures of sheep Cx50 (PDB: 6MHY), Cx46 (PDB: 6MHQ) and designed variants constructed in VMD (grey dotted lines) are overlaid with time-averaged pore profiles obtained during MD production (black lines). Grey shading corresponds to standard error of the mean values obtained during MD (collected every 2 ns, N = 50). Red shading indicates the regions of the NT domain. Asterisk indicates positions of the 9^th^ residue. Pore radii are indicated at the site of the 9^th^ residue obtained from the starting models. **b.** Table summarizing the pore radius at residue 9, the minimum radius (*i.e*., smallest constriction site) and the residue contributing to the minimum radius for each of the constructs. Values obtained from the starting models and following MD equilibration are presented. Values in parenthesis indicate standard deviation (N = 50). Following MD equilibration, all models display a minimum constriction of ~4.6 – 4.8 Å around residue S5, with the exception of Cx50-46NT which displays a significantly wider pore constriction at S5 of ~5.5 Å (P < 0.1 – 0.001) as compared to all other models (over Z-axis values ± 40–46 Å).

**Supplemental Fig. 3.**
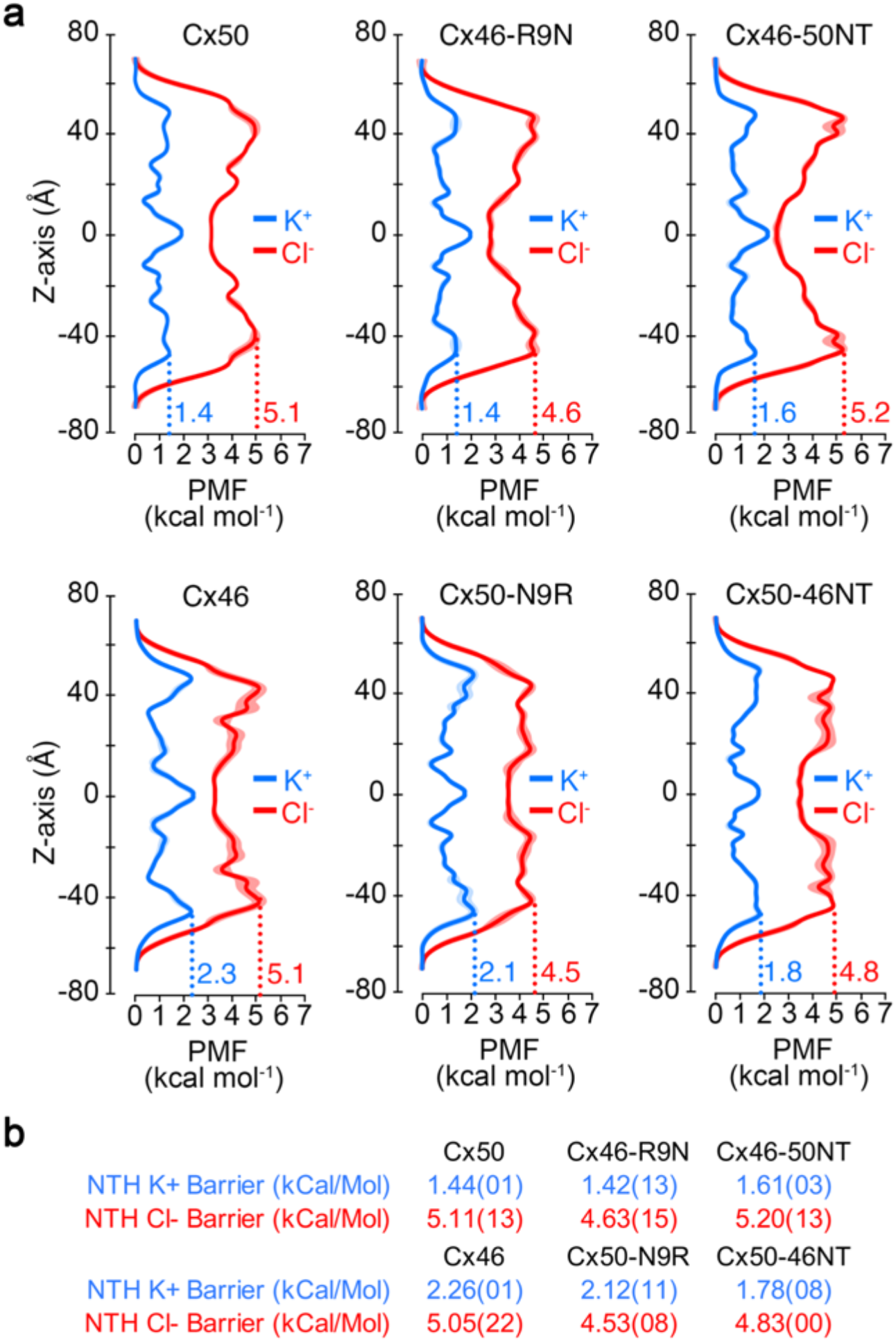
Potential of mean force for ion permeation in Cx50, Cx46 and designed NT domain variants. **a**. Full representation of Potential of Mean Force (PMF) diagrams describing the free-energy landscape (*ΔG*) experienced by K^+^ ions (blue traces) and Cl^-^ ions (red traces) permeating the channel pore. Symmetrized values are shown for acetylated models of Cx50, Cx46-R9N and Cx46-50NT (top row), and Cx46, Cx50-N9R and Cx50-46NT (bottom row), with standard error of the mean (S.E.M.) displayed in lighter shading. Values of energetic barrier within the NT domain are indicated. **b.** Table summarizing the energetic barriers to K^+^ and Cl^-^ ions. Values in parenthesis represent the standard error of the mean (S.E.M.). Note, some variation in the Cl^-^ PMFs is expected due to the larger error associated with the analysis of these rarer anion permeation events.

## References

Aasen T, Mesnil M, Naus CC, Lampe PD & Laird DW. (2016). Gap junctions and cancer: communicating for 50 years. Nat Rev Cancer 16, 775–788.

Abrams CK, Peinado A, Mahmoud R, Bocarsly M, Zhang H, Chang P, Botello-Smith WM, Freidin MM & Luo Y. (2018). Alterations at Arg(76) of human connexin 46, a residue associated with cataract formation, cause loss of gap junction formation but preserve hemichannel function. Am J Physiol Cell Physiol 315, C623–c635.

Bai D, Yue B & Aoyama H. (2018). Crucial motifs and residues in the extracellular loops influence the formation and specificity of connexin docking. Biochim Biophys Acta 1860, 9–21.

Bargiello T & Brink P. (2009). Voltage-gating mechanisms of connexin channels. In Connexins: A Guide, ed. Harris AL & Locke D, pp. 103–128. Humana Press.

Beyer EC, Ebihara L & Berthoud VM. (2013). Connexin mutants and cataracts. Front Pharmacol 4, 43.

Bezanilla F. (2000). The voltage sensor in voltage-dependent ion channels. Physiol Rev 80, 555–592.

Bukauskas FF, Kreuzberg MM, Rackauskas M, Bukauskiene A, Bennett MV, Verselis VK & Willecke K. (2006). Properties of mouse connexin 30.2 and human connexin 31.9 hemichannels: implications for atrioventricular conduction in the heart. Proc Natl Acad Sci U S A 103, 9726–9731.

Bukauskas FF & Verselis VK. (2004). Gap junction channel gating. Biochim Biophys Acta 1662, 42–60.

Delmar M, Laird DW, Naus CC, Nielsen MS, Verselis VK & White TW. (2017). Connexins and Disease. Cold Spring Harbor perspectives in biology.

Flores JA, Haddad BG, Dolan KA, Myers JB, Yoshioka CC, Copperman J, Zuckerman DM & Reichow SL. (2020). Connexin-46/50 in a dynamic lipid environment resolved by CryoEM at 1.9 Å. BioRxiv.

Garcia IE, Prado P, Pupo A, Jara O, Rojas-Gomez D, Mujica P, Flores-Munoz C, Gonzalez-Casanova J, Soto-Riveros C, Pinto BI, Retamal MA, Gonzalez C & Martinez AD. (2016). Connexinopathies: a structural and functional glimpse. BMC Cell Biol 17 Suppl 1, 17.

Gong XQ & Nicholson BJ. (2001). Size selectivity between gap junction channels composed of different connexins. Cell Commun Adhes 8, 187–192.

Goodenough DA & Paul DL. (2009). Gap junctions. Cold Spring Harbor perspectives in biology 1, a002576.

Harris AL, Spray DC & Bennett MV. (1981). Kinetic properties of a voltage-dependent junctional conductance. J Gen Physiol 77, 95–117.

Hopperstad MG, Srinivas M & Spray DC. (2000). Properties of gap junction channels formed by Cx46 alone and in combination with Cx50. Biophys J 79, 1954–1966.

Huang J & MacKerell AD, Jr. (2013). CHARMM36 all-atom additive protein force field: validation based on comparison to NMR data. J Comput Chem 34, 2135–2145.

Humphrey W, Dalke A & Schulten K. (1996). VMD: visual molecular dynamics. J Mol Graph 14, 33-38, 27–38.

Jassim A, Aoyama H, Ye WG, Chen H & Bai D. (2016). Engineered Cx40 variants increased docking and function of heterotypic Cx40/Cx43 gap junction channels. J Mol Cell Card 90, 11–20.

Jurrus E, Engel D, Star K, Monson K, Brandi J, Felberg LE, Brookes DH, Wilson L, Chen J, Liles K, Chun M, Li P, Gohara DW, Dolinsky T, Konecny R, Koes DR, Nielsen JE, Head-Gordon T, Geng W, Krasny R, Wei GW, Holst MJ, McCammon JA & Baker NA. (2018). Improvements to the APBS biomolecular solvation software suite. Protein Sci 27, 112–128.

Kanaporis G, Mese G, Valiuniene L, White TW, Brink PR & Valiunas V. (2008). Gap junction channels exhibit connexin-specific permeability to cyclic nucleotides. J Gen Physiol 131, 293–305.

Kim MS, Gloor GB & Bai D. (2013). The distribution and functional properties of Pelizaeus-Merzbacher-like disease-linked Cx47 mutations on Cx47/Cx47 homotypic and Cx47/Cx43 heterotypic gap junctions. The Biochemical journal 452, 249–258.

Koval M, Molina SA & Burt JM. (2014). Mix and match: investigating heteromeric and heterotypic gap junction channels in model systems and native tissues. FEBS Lett 588, 1193–1204.

Kronengold J, Srinivas M & Verselis VK. (2012). The N-terminal half of the connexin protein contains the core elements of the pore and voltage gates. The Journal of membrane biology 245, 453–463.

Mackay DS, Bennett TM, Culican SM & Shiels A. (2014). Exome sequencing identifies novel and recurrent mutations in GJA8 and CRYGD associated with inherited cataract. Hum Genomics 8, 19.

Maeda S, Nakagawa S, Suga M, Yamashita E, Oshima A, Fujiyoshi Y & Tsukihara T. (2009). Structure of the connexin 26 gap junction channel at 3.5 A resolution. Nature 458, 597–602.

Mathias RT, White TW & Gong X. (2010). Lens gap junctions in growth, differentiation, and homeostasis. Physiol Rev 90, 179–206.

Micheal S, Niewold ITG, Siddiqui SN, Zafar SN, Khan MI & Bergen AAB. (2018). Delineation of Novel Autosomal Recessive Mutation in GJA3 and Autosomal Dominant Mutations in GJA8 in Pakistani Congenital Cataract Families. Genes 9.

Moreno AP, Berthoud VM, Perez-Palacios G & Perez-Armendariz EM. (2005). Biophysical evidence that connexin-36 forms functional gap junction channels between pancreatic mouse beta-cells. Am J Physiol Endocrinol Metab 288, E948–956.

Musa H, Fenn E, Crye M, Gemel J, Beyer EC & Veenstra RD. (2004). Amino terminal glutamate residues confer spermine sensitivity and affect voltage gating and channel conductance of rat connexin40 gap junctions. J Physiol 557, 863–878.

Myers JB, Haddad BG, O’Neill SE, Chorev DS, Yoshioka CC, Robinson CV, Zuckerman DM & Reichow SL. (2018). Structure of native lens connexin 46/50 intercellular channels by cryo-EM. Nature 564, 372–377.

Oh S, Rubin JB, Bennett MV, Verselis VK & Bargiello TA. (1999). Molecular determinants of electrical rectification of single channel conductance in gap junctions formed by connexins 26 and 32. J Gen Physiol 114, 339–364.

Pal JD, Liu X, Mackay D, Shiels A, Berthoud VM, Beyer EC & Ebihara L. (2000). Connexin46 mutations linked to congenital cataract show loss of gap junction channel function. Am J Physiol Cell Physiol 279, C596–602.

Paul DL, Ebihara L, Takemoto LJ, Swenson KI & Goodenough DA. (1991). Connexin46, a novel lens gap junction protein, induces voltage-gated currents in nonjunctional plasma membrane of Xenopus oocytes. J Cell Biol 115, 1077–1089.

Pettersen EF, Goddard TD, Huang CC, Couch GS, Greenblatt DM, Meng EC & Ferrin TE. (2004). UCSF Chimera--a visualization system for exploratory research and analysis. J Comput Chem 25, 1605–1612.

Phillips JC, Braun R, Wang W, Gumbart J, Tajkhorshid E, Villa E, Chipot C, Skeel RD, Kale L & Schulten K. (2005). Scalable molecular dynamics with NAMD. J Comput Chem 26, 1781–1802.

Purnick PE, Oh S, Abrams CK, Verselis VK & Bargiello TA. (2000). Reversal of the gating polarity of gap junctions by negative charge substitutions in the N-terminus of connexin 32. Biophys J 79, 2403–2415.

Rubinos C, Villone K, Mhaske PV, White TW & Srinivas M. (2014). Functional effects of Cx50 mutations associated with congenital cataracts. Am J Physiol Cell Physiol 306, C212–220.

Saez JC, Berthoud VM, Branes MC, Martinez AD & Beyer EC. (2003). Plasma membrane channels formed by connexins: their regulation and functions. Physiol Rev 83, 1359–1400.

Sakai R, Elfgang C, Vogel R, Willecke K & Weingart R. (2003). The electrical behaviour of rat connexin46 gap junction channels expressed in transfected HeLa cells. Pflugers Arch 446, 714–727.

Shearer D, Ens W, Standing K & Valdimarsson G. (2008). Posttranslational modifications in lens fiber connexins identified by off-line-HPLC MALDI-quadrupole time-of-flight mass spectrometry. Invest Ophthalmol Vis Sci 49, 1553–1562.

Smart OS, Neduvelil JG, Wang X, Wallace BA & Sansom MS. (1996). HOLE: a program for the analysis of the pore dimensions of ion channel structural models. J Mol Graph 14, 354–360, 376.

Sohl G & Willecke K. (2004). Gap junctions and the connexin protein family. Cardiovasc Res 62, 228–232.

Sosinsky GE & Nicholson BJ. (2005). Structural organization of gap junction channels. Biochim Biophys Acta 1711, 99–125.

Spray DC, Harris AL & Bennett MV. (1981). Equilibrium properties of a voltage-dependent junctional conductance. Journal of General Physiology 77, 77–93.

Srinivas M, Costa M, Gao Y, Fort A, Fishman GI & Spray DC. (1999). Voltage dependence of macroscopic and unitary currents of gap junction channels formed by mouse connexin50 expressed in rat neuroblastoma cells. J Physiol 517 (Pt 3), 673–689.

Sukhareva M, Hackos DH & Swartz KJ. (2003). Constitutive activation of the Shaker Kv channel. J Gen Physiol 122, 541–556.

Sun Y, Yang YQ, Gong XQ, Wang XH, Li RG, Tan HW, Liu X, Fang WY & Bai D. (2013). Novel germline GJA5/connexin40 mutations associated with lone atrial fibrillation impair gap junctional intercellular communication. Human mutation 34, 603–609.

Tong JJ & Ebihara L. (2006). Structural determinants for the differences in voltage gating of chicken Cx56 and Cx45.6 gap-junctional hemichannels. Biophysical Journal 91, 2142–2154.

Tong JJ, Liu X, Dong L & Ebihara L. (2004). Exchange of gating properties between rat cx46 and chicken cx45.6. Biophys J 87, 2397–2406.

Tong X, Aoyama H, Tsukihara T & Bai D. (2014). Charge at the 46th residue of connexin50 is crucial for the gap-junctional unitary conductance and transjunctional voltage-dependent gating. J Physiol 592, 5187–5202.

Trexler EB, Bennett MV, Bargiello TA & Verselis VK. (1996). Voltage gating and permeation in a gap junction hemichannel. Proc Natl Acad Sci U S A 93, 5836–5841.

Trexler EB, Bukauskas FF, Kronengold J, Bargiello TA & Verselis VK. (2000). The first extracellular loop domain is a major determinant of charge selectivity in connexin46 channels. Biophys J 79, 3036–3051.

Veenstra RD, Wang HZ, Beyer EC, Ramanan SV & Brink PR. (1994). Connexin37 forms high conductance gap junction channels with subconductance state activity and selective dye and ionic permeabilities. Biophysical Journal 66, 1915–1928.

Vernon RM, Chong PA, Tsang B, Kim TH, Bah A, Farber P, Lin H & Forman-Kay JD. (2018). Pi-Pi contacts are an overlooked protein feature relevant to phase separation. eLife 7.

Verselis VK, Ginter CS & Bargiello TA. (1994). Opposite voltage gating polarities of two closely related connexins. Nature 368, 348–351.

Wang Z & Schey KL. (2009). Phosphorylation and truncation sites of bovine lens connexin 46 and connexin 50. Exp Eye Res 89, 898–904.

White TW, Bruzzone R, Goodenough DA & Paul DL. (1994a). Voltage gating of connexins. Nature 371, 208–209.

White TW, Bruzzone R, Wolfram S, Paul DL & Goodenough DA. (1994b). Selective interactions among the multiple connexin proteins expressed in the vertebrate lens: the second extracellular domain is a determinant of compatibility between connexins. J Cell Biol 125, 879–892.

Xin L, Gong XQ & Bai D. (2010). The role of amino terminus of mouse Cx50 in determining transjunctional voltage-dependent gating and unitary conductance. Biophys J 99, 2077–2086.

Xin L, Nakagawa S, Tsukihara T & Bai D. (2012a). Aspartic acid residue D3 critically determines Cx50 gap junction channel transjunctional voltage-dependent gating and unitary conductance. Biophys J 102, 1022–1031.

Xin L, Sun Y & Bai D. (2012b). Heterotypic connexin50/connexin50 mutant gap junction channels reveal interactions between two hemichannels during transjunctional voltage-dependent gating. J Physiol 590, 5037–5052.

Ye Y, Wu M, Qiao Y, Xie T, Yu Y & Yao K. (2019). Identification and preliminary functional analysis of two novel congenital cataract associated mutations of Cx46 and Cx50. Ophthalmic genetics 40, 428–435.

Yifrach O & MacKinnon R. (2002). Energetics of pore opening in a voltage-gated K(+) channel. Cell 111, 231–239.

Zhang L, Liang Y, Zhou Y, Zeng H, Jia S & Shi J. (2018). A Missense Mutation in GJA8 Encoding Connexin 50 in a Chinese Pedigree with Autosomal Dominant Congenital Cataract. Tohoku J Exp Med 244, 105–111.

